# Converging roles of glutamate receptors in domestication and prosociality

**DOI:** 10.1101/439869

**Authors:** Thomas O’Rourke, Cedric Boeckx

## Abstract

The present paper highlights the prevalence of signals of positive selection on genes coding for glutamate receptors—most notably kainate and metabotropic receptors—in domesticated animals and anatomically modern humans. Relying on their expression in the central nervous system and phenotypes associated with mutations in these genes, we claim that regulatory changes in kainate and metabotropic receptor genes have led to alterations in limbic function and Hypothalamic-Pituitary-Adrenal axis regulation, with potential implications for the emergence of unique social behaviors and communicative abilities in (self-)domesticated species.

## 1 Introduction

Under one account of recent human evolution, selective pressures on prosocial behaviors led not only to a species-wide reduction in reactive aggression and the extension of our social interactions [1, 2], but also left discernible physical markers on the modern human phenotype, including our characteristically “gracile” anatomy [3, 4].

It has long been noted that these morphological differences resemble those of domesticated species when compared with their wild counterparts [5]. Experimental observation of domestication unfolding in wild farm-bred silver foxes has unequivocally shown that selection for tameness alone can affect developmental trajectories to bring about a suite of physiological and behavioral traits indicative of the “domestication syndrome”[6]. This raises the possibility that morphological changes in *Homo sapiens* resulted from selective pressures on reduced reactivity to encounters with conspecifics. In turn, this predicts overlapping regions of selection and convergent physiological effects in the genomes of domesticated species and modern humans.

In a recent comparative study, we have shown that genes with signals of positive selection pooled across different domesticated species — dog, cattle, cat, and horse — have above-chance overlap with genes exhibiting signals of selection in AMH, suggestive of convergent evolutionary processes [4]. Signals on glutamate receptor genes were identified more consistently than any other gene class across human and domesticate selective-sweep studies, and will be the focus of this paper.

Glutamate is the primary excitatory neurotransmitter in the vertebrate nervous system, essential for fast synaptic transmission and plasticity, learning, memory, and modulation of Hypothalamic-Pituitary-Adrenal (HPA) activity. [7]. The 26 glutamate receptors are primarily localized at synaptic nerve terminals in the brain and are divisible into two broad families (ionotropic and metabotropic). There are various widely used names for each receptor and corresponding gene in the literature. For ease of reference, see Table 1.

**Table 1:** Glutamate receptors. (Throughout the present study we use HUGO nomenclature to refer to both receptor genes and proteins.)

| Ionotropic |  |  |  |  |  |  |  |  |
| --- | --- | --- | --- | --- | --- | --- | --- | --- |
| NMDA |  |  | AMPA |  |  | Kainate |  |  |
| HUGO | Aliases |  | HUGO | Aliases |  | HUGO | Aliases |  |
| GRIN1 | NR1 | GluN1 | GRIA1 | GLUR1 | GLUA1 | GRIK1 | GLUR5 | GLUK1 |
| GRIN2A | NR2A | GluN2A | GRIA2 | GLUR2 | GLUA2 | GRIK2 | GLUR6 | GLUK2 |
| GRIN2B | NR2B | GluN2B | GRIA3 | GLUR3 | GLUA3 | GRIK3 | GLUR7 | GLUK3 |
| GRIN2C | NR2C | GluN2C | GRIA4 | GLUR4 | GLUA4 | GRIK4 | KA1 | GLUK4 |
| GRIN2D | NR2D | GluN2D |  |  |  | GRIK5 | KA2 | GLUK5 |
| GRIN3A | NR3A | GluN3A |  |  |  | Delta |  |  |
| GRIN3B | NR3B | GluN3B |  |  |  | GRID1 | GLUD1 |  |
|  |  |  |  |  |  | GRID2 | GLUD2 |  |

| Metabotropic |  |  |  |  |  |  |  |  |
| --- | --- | --- | --- | --- | --- | --- | --- | --- |
| Group I |  |  | Group II |  |  | Group III |  |  |
| HUGO | Aliases |  | HUGO | Aliases |  | HUGO | Aliases |  |
| GRM1 | mGluR1 | mGlu1 | GRM2 | mGluR2 | mGlu2 | GRM4 | mGluR4 | mGlu4 |
| GRM5 | mGluR5 | mGlu5 | GRM3 | mGluR3 | mGlu3 | GRM6 | mGluR6 | mGlu6 |
|  |  |  |  |  |  | GRM7 | mGluR7 | mGlu7 |
|  |  |  |  |  |  | GRM8 | mGluR8 | mGlu8 |

Given their importance for the normal functioning of the organism, glutamate receptors are rarely subject to extensive structural changes from one species to the next [8, 9]. Despite this high structural conservation, there are significant differences between humans and chimpanzees in the cortical expression of glutamate receptor genes [10, 11], suggesting that changes to regulatory regions may have had important functional consequences for the emergence of the human cognitive phenotype. Similarly, the vast majority of selective sweeps or high-frequency changes on glutamate receptor genes in AMH relative to archaic Homo are found in regulatory regions that control gene expression [12, 13].

Like the tameness of domesticates towards carers, prosociality among humans necessitates a reduction in fearful and aggressive reactions to encounters with conspecifics. Fear, anxiety, and aggression are stress responses, mediated across vertebrates by the hypothalamic-pituitary-adrenal (HPA) axis [14, 15], the hypofunction of which has been proposed to be a key mechanism in the development of tame behaviors in domesticates [16, 17]. Here we articulate the hypothesis that increased modulatory actions of glutamate receptors have brought about attenuation of the HPA stress response in ours and domesticated species. We first characterize the extent to which certain (sub)families of glutamate receptor genes exhibit shared signals of selection in domesticates and modern humans, comparing these with genes for other neurotransmitter and nuclear hormone receptors. We then discuss the likely functional implications of these changes with reference to gene expression data and evidence from developmental and psychiatric disorders.

## 2 Results

In earlier work, we have pointed out the intersection of glutamate receptor genes showing signals of selection in humans, dogs, cats, cattle, and horses [4]. Here we take into account a much broader range of human and domestication studies showing changes in AMH, dogs, cats, cattle, horses, foxes, sheep, pigs, rabbits, yaks, goats, guinea pigs, chickens, and ducks. We compare changes on the 26 glutamate receptor genes with those for genes encoding 462 other receptors from equivalent classes (G-protein coupled receptors and ligand-gated ion channels), as well the other major receptor superfamilies in the central nervous system with potential relevance to domestication events (receptor kinases and nuclear hormone receptors; see Supplementary Table S1)

Although we focus here on signals of selection on glutamate receptor genes, changes to other glutamatergic signaling genes have also been identified in modern human and domestication studies. These include glutamate transporter, accessory subunit, and related G-protein signaling cascade genes. We review some of the most noteworthy of these in Supplementary Section S1.

At least one glutamate receptor gene shows signals of selection in all of the species in Table 2. Overlapping signals of selection were most consistently detected on kainate and Group II and III metabotropic receptor genes across modern human and animal domestication studies (see Table 2 and Figure 1).

**Table 2:** Signals of selection, high-frequency changes, introgression, and differential expression of glutamate receptor genes in AMH and domesticated species.

|  | AMH<br>[12, 13, 18, 19, 20, 21] | Dog<br>[22, 23, 24, 25, 26] | Cattle<br>[27] | Cat<br>[28] | Horse<br>[29] | Fox<br>[30, 31] | Yak<br>[32, 33] | Sheep<br>[34] | Goat<br>[35, 36, 37] | Pig<br>[38, 39, 40] | Rabbit<br>[41] | Guinea Pig<br>[42] | Chicken<br>[43] | Duck<br>[44] |
| --- | --- | --- | --- | --- | --- | --- | --- | --- | --- | --- | --- | --- | --- | --- |
| GRIN1 |  |  |  |  |  |  |  |  |  |  |  |  |  |  |
| GRIN2A |  |  |  |  |  |  | ▲ |  |  |  |  |  |  |  |
| GRIN2B |  |  |  |  |  | ■ | ▲ |  |  |  |  |  |  |  |
| GRIN2C | ■ |  |  |  |  |  |  |  |  |  |  |  |  |  |
| GRIN2D | ● |  |  |  |  | ◆ |  |  |  |  |  |  |  |  |
| GRIN3A | ● |  |  |  |  |  | ▲ |  |  |  |  |  |  | ● |
| GRIN3B |  |  |  |  |  |  |  |  |  |  |  |  |  |  |
| GRIA1 |  |  |  | ● |  |  |  |  |  |  |  |  |  |  |
| GRIA2 | ● |  |  | ● |  |  |  |  |  | ● |  |  |  |  |
| GRIA3 |  |  |  |  |  |  |  |  |  |  |  |  |  |  |
| GRIA4 |  |  |  |  |  |  | ▲ |  |  |  |  |  |  |  |
| GRIK1 |  |  |  |  |  |  |  |  |  |  |  |  |  |  |
| GRIK2 |  | ● ◆ |  |  |  |  | ▲ |  |  |  | ● |  |  | ● |
| GRIK3 | ● | ● ● ● ● ● | ● |  |  |  | ▲ | ● |  |  |  |  |  |  |
| GRIK4 | ■ |  |  |  |  |  |  |  |  |  |  |  |  |  |
| GRIK5 | ● ● ■ |  |  |  |  |  |  |  |  |  |  |  |  |  |
| GRM1 |  |  |  |  |  |  |  |  |  |  |  |  | ● |  |
| GRM2 | ● ● ■ |  |  |  |  |  |  |  |  |  |  |  |  |  |
| GRM3 | ● ● |  |  |  |  | ■ |  |  |  |  |  |  |  |  |
| GRM4 |  |  |  |  |  |  | ▲ |  |  |  |  | ◆ |  |  |
| GRM5 |  |  |  |  |  |  |  |  |  |  |  |  |  |  |
| GRM6 | ■ |  |  |  |  | ■ |  |  |  |  |  |  |  |  |
| GRM7 | ■ |  |  |  |  |  |  |  | ● | ● |  |  |  |  |
| GRM8 | ● | ● |  |  |  |  |  | ● |  | ● ● |  |  |  |  |
| GRID1 |  |  |  |  | ● |  |  |  |  |  |  |  |  |  |
| GRID2 | ● |  |  |  |  |  |  |  |  |  |  |  |  |  |

- Selective sweep study identifying differences between AMH and archaics or domesticates and their wild ancestors - High frequency changes differentiating AMH and archaics or domesticates and their wild ancestors - ◆ Differential brain expression between domesticated species and wild ancestors - ▲ Introgression study

**Figure 1:**
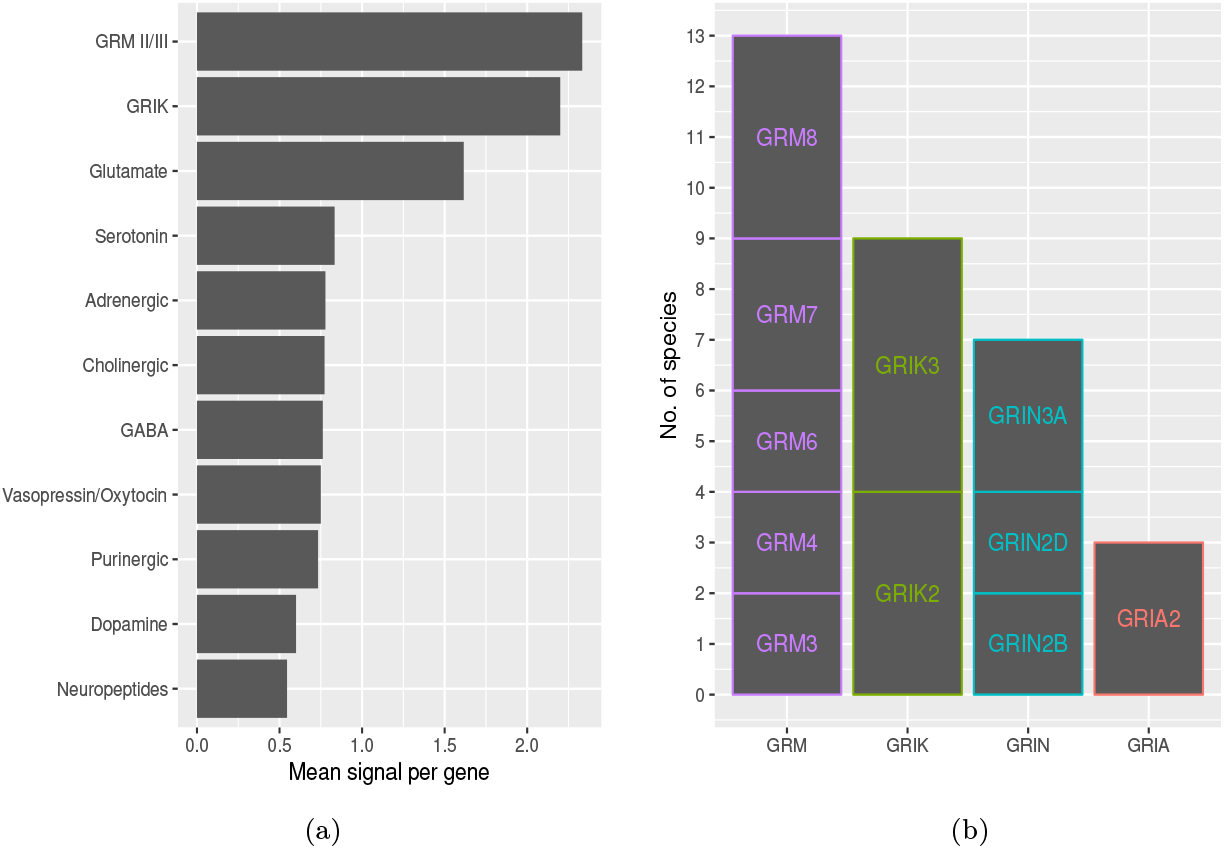
Signals of selection, introgression, high-frequency changes, or expression differences on abundant neurotransmitter receptor and glutamate receptor genes, plotted by gene group. For full comparisons of receptor types, see supplementary Table S1 (a) : Comparison of mean signals across major receptor types (b) : Number of species for which signals are detected in at least two species per glutamate receptor gene.

Out of the thirty-two instances where differences were detected on functioning ionotropic glutamate receptor genes (NMDA, AMPA, or kainate), eighteen were detected among the kainate receptor genes, and fourteen of these occurred either on *GRIK2* or *GRIK3. GRIK3* exhibits signals of selection in modern humans, dogs, cattle, sheep, and of introgression in yaks, while *GRIK2* shows signals of selection in dogs (accompanied by increased brain expression as compared with wolves), rabbits, and ducks, and of introgression in yaks.

Metabotropic receptors are the other major subclass of glutamate receptor genes that display consistently convergent signals among domesticated species and modern humans, with members of Group II (*GRM2* and *GRM3*) and, in particular, Group III (*GRM4, GRM6, GRM7*, and *GRM8*) exhibiting signals of selection across domesticate and human selective-sweep studies. Signals on *GRM8* have been detected in dogs, sheep, pigs, and humans. Relative to Neanderthals and Denisovans, modern humans show signals of selection on *GRM2* and *GRM3* (this latter gene detected on both the Yoruba and KhoeSan branches of our lineage).

Eighteen out of the nineteen instances where signals of selection, expression differences, high-frequency missense changes, or introgression were detected on metabotropic glutamate receptor genes, these occurred in Group II or Group III subfamilies. The only exception to this pattern is the detection of a selective sweep on *GRM1* in the chicken. Group II and III metabotropic receptors share structural and functional similarities, which differentiate them from Group I receptors. Most strikingly, Group II and III receptor subunits can form functional heterodimers with each other across subfamilies, while they are unable to do so with Group I subunits [45]. Moreover, activation of Group II and III receptors primarily inhibits adenylyl-cyclase signaling, whereas Group I receptors potentiate this signaling cascade [46]. These facts suggest that Group II and III metabotropic receptors form a class that is mechanistically distinct from Group I. This, in turn, raises the possibility that signals of selection across these two subfamilies may be instances of convergent evolutionary processes.

For the comparison between the 488 receptor genes and their relevant (sub) families, we considered a gene to potentially be implicated in a domestication event or in modern human evolution (to exhibit a ‘signal’) if it was identified in at least one of the studies in Table 2. Detection of a single gene across different studies of a single species were counted as a one signal, leaving a total of 401 putative signals detected across the 488 genes (a receptor-wide mean of 0.82 signals per gene). Receptor genes were subdivided into 106 distinct multigene families (or 114 when subfamily divisions were included).

Group III metabotropic receptor genes exhibited signals at a rate higher than those of any other multigene receptor (sub)families (mean of 2.75 signals per gene), including receptor genes for the most abundant neurotransmitters in the brain, such as GABA (mean: 0.76; GABA_A_ Beta mean: 1.33; GABA_B_ mean: 1.5), serotonin (mean: 0.83; HTR2 family: 1.33), dopamine (mean: 0.6), (nor)epinephrine (mean: 0.78), or acetylcholine receptors (mean: 0.77). In general, signals on these receptor gene families tended to track the neurotransmitter-wide average of 0.82. Signals on kainate receptor genes occurred at a rate considerably higher than any of the above (sub)families (mean: 2.2). Of the 114 multigene (sub)families examined, only corticosteroid receptors (mean: 2.5), and Adhesion G protein-coupled receptor (ADGR) subfamilies B (mean 2.33) and D (mean 2.5) exhibited signals at higher rates (see Table S1).

Both the corticosteroid and ADGRD families contain only two receptor genes, meaning that each family had a total of just five signals across the thirty domestication and modern human evolution studies examined here. By contrast, *GRIK3* alone exhibited five signals of selection across these studies, and *GRIK2* four. The small size of the corticosteroid and ADGRD receptor families may well have contributed to the high mean signals highlighted here. To control for this, we carried out a Monte Carlo simulation to determine the chance likelihood that each of the receptor (sub)families under comparison here should the preponderance of signals that they do, given the family size. This analysis showed that the prevalence of signals occurring on metabotropic family III or kainate receptor (sub)families was unlikely to occur by chance (p<0.00001 and p=0.037, respectively). By contrast, the chance likelihood of signals occurring at the rate they do on receptor families with comparable means (ADGRB, ADGRD, and corticosteroid) was considerably higher (p=0.12, p=0.265, and p=0.265, respectively). Given their ability to form functioning heterodimers, Group II and III metabotropic receptors could be considered a single functional subfamily. The fourteen signals occurring across these six genes was highly unlikely to have occurred by chance, given the receptor-wide average (p = 0.003)

One-way ANOVA comparisons were carried out among all multigene receptor (sub)families identified here in order to determine if subfamilies with the highest mean signals showed significant divergence from those with lower means. Out of the twenty-two pairwise comparisons that showed significant differences between means, twenty involved metabotropic subfamily III, while the two other significant comparisons involved the kainate receptor family (see Table S2). When one considers that alterations to neurotransmission are highlighted in many of the domestication studies analysed here, these findings suggest that signals on metabotrobic III and kainate receptor (sub)families are, at least, significantly higher than background genome-wide signals of selection under domestication.

Averaging across multigene families allows for the identification of convergent signals on distinct genes with similar functional properties. However, this does not allow for analysis of single-gene receptor families, and it may also obscure above-chance convergent signals on individual genes within multigene families. *GRIK3* was one of only two genes exhibiting changes in five species, the other being *ADGRB3* (Likelihood according to Monte Carlo simulation: p=0.004). The ADGRB3 subunit (also known as BAI3), like kainate receptors, is bound by C1ql proteins, modulates excitatory transmission, as per kainate and metabotropic glutamate receptors, and is implicated in similar neurodevel-opmental and stress disorders as genes from these (sub) families [47, 48, 49].

*GRIK2* and *GRM8* were two of only five genes that were identified in four domesticates (including modern humans in the case of *GRM8*) (Likelihood according to Monte Carlo simulation: p=0.043). *KIT*, a gene involved in the differentiation of neural crest precursor cells into melanocytes, and important in selection for coat color in domesticated species [17], was also identified in four different species. Three glutamate receptor genes (*GRM7, GRIN3A*, and *GRIA2*) were identified among the ten genes with changes detected in three species. Functional enrichment (Gene Ontology) analyses of these seventeen genes with signals in three or more species showed glutamate receptor signaling to be the most highly-enriched pathway in the Biological Process category, with glutamatergic activity being in three of the top five pathways highlighted under Molecular Function. An extended functional analysis of the 92 genes showing signals of selection in two or more species highlighted ‘Glutamatergic synapse’ as the second most prominent KEGG pathway, behind the more general (and subsuming) category ‘Neuroactive ligand-receptor interaction’.

In several of the studies cited in Table 2, signals of selection on glutamate receptor genes have been suggested as potentially important for behavioral changes during the domestication process [23, 27, 28, 29, 30, 39, 44]. However, such suggestions often form part of broader discussion on changes to genes related to the central nervous system (CNS) under domestication, and no mech anistic details are provided. Furthermore, discussion is usually limited to highlighting the potential importance of glutamatergic-signaling changes in learning, memory, or excitatory transmission in a single domesticated species under study, and tends to focus on cortical regions.

To our knowledge, no study to date has sought to explore the extent to which glutamate receptor genes show signals of selection across a large number of domesticated species. More significantly, we are unaware of any previous study that has identified kainate and metabotropic receptor genes as the most prevalent targets of selective sweeps under domestication and in recent human evolution. The consistency with which these receptor genes are identified across domestication studies makes them prime candidates for being involved in the emergence of tameness.

If an argument is to be made for the convergent involvement of kainate and metabotropic glutamate receptor genes in domesticated tameness and human prosociality, evidence should point to the shared participation of these receptors in modulating the stress response across these species. Table 3 summarizes evidence from human, model organism, and animal gene-behavior-correlation studies that kainate and Group II and III metabotropic receptors highlighted here are implicated in developmental, neuropsychiatric, stress, and mood disorders, as well as divergent tame and agonistic phenotypes in non-human species. (Detailed discussion of these studies can be found in Supplementary section S2.)

**Table 3:** Human and Domesticated phenotypes associated with kainate and metabotropic receptor genes. Detailed discussion of these associations, including references can be found in Supplementary section S2. DD/ID - Developmental Delay/Intellectual Disability, ADHD - Attention Deficit Hyperactivity Disorder, ASD - Autism Spectrum Disorder, SCZ - Schizophrenia, BPD - Bipolar Disorder, OCD - Obsessive Compulsive Disorder, ANX - Anxiety Disorder, MDD - Major Depression, SR/Fear - Startle Response/Fear.

| GENE | Associated Human phenotypes |  |  |  |  |  |  |  |  | Tame and aggressive animal phenotypes |
| --- | --- | --- | --- | --- | --- | --- | --- | --- | --- | --- |
|  | DD/ID | ADHD | ASD | SCZ | BPD | OCD | ANX | MDD | SR/Fear |  |
| GRIK2 | * | * | * | * | * | * | * | * | * | Retention of dog-like polymorphisms on GRIK2 in tame Czechoslovakian Wolf-Dog hybrid [58] |
| GRIK3 | * |  | * | * | * |  | * | * | * | Signals of selection on GRIK3 differentiating cattle with agonistic and tame behaviors [59] |
| GRIK5 |  |  | * | * | * |  | * |  |  |  |
| GRM2 |  | * | * | * |  | * | * | * | * |  |
| GRM3 |  | * | * | * | * | * | * | * | * | Fixed missense mutation on GRM3 differentiating aggressive from tame foxes [30] |
| GRM4 |  | * | * | * | * |  | * | * | * |  |
| GRM7 |  | * | * | * | * | * | * | * | * | Lower nucleotide variability on GRM7 in docile Apennine Brown Bear [60] |
| GRM8 | * | * | * | * |  |  | * | * | * | Lower nucleotide variability on GRM8 in docile Apennine Brown Bear [60] |

Note: Widespread use of non-selective Group II agonists that act on both GRM2 and GRM3 subunits in model organisms makes it difficult to always separate subunit associations with disorders. Where a disorder has been associated with Group II receptors, an asterisk is placed in the row for both GRM2 and GRM3.

Schizophrenia is among the disorders most regularly associated with mutations on these metabotropic and kainate receptor genes. In humans, prenatal stress is a risk factor for the development of schizophrenia in adult offspring [50, 51]. Pharmacological agonists of Group II metabotropic receptors reduce schizophrenia-like phenotypes in adult offspring of prenatally stressed mice [52]. Furthermore, high glucocorticoid inputs to the hippocampus reduce the expression of kainate receptors, and schizophrenics have been found express significantly reduced kainate receptors in this region [53, 54]. More broadly, heightened stress experienced during pregnancy can lead to a “persistently hyperactive” HPA axis in offspring, increasing children’s propensity to develop Attention Deficit Hyperactivity Disorder (ADHD), as well as adult anxiety and reactivity to stress, while in rats, prenatal stress decreases the propensity to play in juvenile offspring and impairs sociality and extinction of conditioned fear lasting into adulthood [55, 56, 57].

Prenatal stress can, then, contribute to the emergence of the same neurodevelopmental, neuropsychiatric, stress, and mood disorders commonly associated with altered expression of metabotropic and kainate receptor genes. We hypothesize that selective sweeps on these genes are markers of convergent positive selection on an attenuated stress response in both ancient modern humans and early domesticated species. We propose that enhanced prenatal modulation by these receptors of stress responses to human contact in (pre-)domesticated and archaic human females provided an important first step in the emergence tameness and prosociality. In section 3, we explore the neurobiological evidence for this proposal, focusing on the roles of kainate and metabotropic glutamate receptors in the stress-response cascade.

## 3 Discussion

Alterations to the HPA axis are considered to be essential for the emergence of tameness in different (indeed competing) theories of domestication [17, 61]. Glutamatergic signaling acts as a prominent regulator of HPA activity and has been identified among the top enriched pathways across studies of aggression [7, 62, 63, 64]. The association of kainate and Group II/III metabotropic receptor genes with multiple stress disorders implicates them in altered HPA-axis activity. Given the predominantly modulatory, as opposed to excitatory, functions of both kainate and Group II and III metabotropic receptors [46, 65], we argue that selective sweeps on their respective genes are markers of decreased HPA reactivity in humans and (pre-)domesticated species.

Below, we propose a mechanism for how these receptor subfamilies modulate glutamatergic signaling to alter developmental trajectories and, subsequently, the HPA stress response in both humans and domesticated species. Our argument relies on three pieces of evidence: First, that alterations to the HPA axis are common across domesticated species versus their wild counterparts, and in non-reactive versus reactively aggressive humans; second, evidence that the genes highlighted here are extensively expressed in limbic and hypothalamic brain regions crucial for controlling the stress response; and third, evidence that disturbance of this expression alters the stress response.

### Alterations to the stress response in domesticated species and modern humans

In response to stress, corticotrophin releasing hormone (CRH) is synthesized in the paraventricular nucleus (PVN) of the hypothalamus. This induces adreno-corticotrophin (ACTH) release from the anterior pituitary gland, which, in turn, stimulates the release of glucocorticoids (GCs: primarily cortisol and corticosterone) from the adrenal gland [66]. GCs are “the principal end-products of the HPA axis”, which help to maintain homeostatic balance in the organism [67]. They also provide feedback directly to neurons in the PVN [14, 68], or via other brain regions, particularly limbic structures, including the hippocampus, thereby modulating CRH release and the HPA stress response [69, 67]. Thus, GC measures can be an accurate indicator of stress response in vertebrates, once basal and stress-response measures can be differentiated [70].

Domesticated foxes, sheep, bengalese finches, and ducks have lower basal GC levels than their wild ancestors or other closely related wild comparators [71, 72, 73, 74, 75, 76, 77]. In the duck and the fox, differences are particularly marked in prenatal and juvenile development, respectively [76, 77, 71]. Compared to their wild ancestors, Guinea pigs and chickens have a lower spike in GCs in response to stress [78, 79]. Although there is no extant ancestral comparator of neuroendocrine function in AMH, our species has considerably lower basal plasma cortisol levels than chimpanzees and most other primates [80].

Within the human population, variability in GC levels correlate with different individual stress responses, which mirrors findings in laboratory rats. Acute GC increases accompany bouts of reactive aggression, while chronically high basal levels have been found to correlate with increased anxiety and major depression, and may be implicated in reduced aggressive tendencies [15]. Chronically low GC levels can correlate with antisocial personality disorder, callous, unemotional tendencies, and externalizing behaviors in children, as well as aggressive delinquency in adults. Proactively aggressive or non-aggressive children tend to have a lower spike in GC levels in response to frustrating tasks than reactively aggressive children [81]. Psychopathic adults (who often exhibit pathological proactive aggression) tend to have no cortisol reactivity to frustrating tasks [15].

The above studies suggest that, from early development into adulthood, lower basal GC levels are shared by domesticates and modern humans relative to closely related extant wild species. Moreover decreased GC spikes in response to stress are common to domesticates and non-aggressive or proactively aggressive modern humans versus reactively aggressive individuals. These findings are consistent with the view that prosocial selective pressures have led to a reduction in reactive over and above proactive aggression in recent human evolution [1]. It could be considered that proactive aggressors within modern human populations exhibit a pathological version of the non-reactive phenotype that has been under positive selection and is associated with HPA-axis hypofunction under stress.

### Kainate and metabotropic receptor expression in brain regions crucial for HPA regulation

The HPA axis is centrally regulated by the limbic system, primarily through amygdalar processing of perceptual inputs, which are relayed via the bed nucleus of the stria terminalis (BNST) to the paraventricular nucleus (PVN) in the hypothalamus. The limbic system also mediates feedback mechanisms, whereby glucocorticoids and mineralocorticoids act upon receptors in the hippocampus and medial prefrontal cortex, which connect to the PVN via the BNST and lateral septum. Feedback also occurs directly on cells in the PVN to modulate HPA reactivity. Feedforward mechanisms, further potentiating the stress response, are relayed from the amygdala to the PVN via the BNST. [82, 69, 68].

Glutamatergic and GABAergic signaling are the central mediators of each of these aspects of HPA (re)activity, and the kainate and metabotropic receptor subfamilies discussed here play prominent roles in modulating release of both neurotransmitters. These receptors are extensively expressed in limbic regions crucial for modulating the stress response. Figure 2 highlights these expression patterns, including both Group II and III metabotropic receptors, given their shared structural and functional properties. A detailed overview of what is known about kainate and Group II and III metabotropic receptor expression in the developing and adult brain can be found in Supplementary section S3. In the subsection that follows, we propose a mechanism by which metabotropic and kainate receptors modulate, and are modulated by, HPA activity.

**Figure 2:**
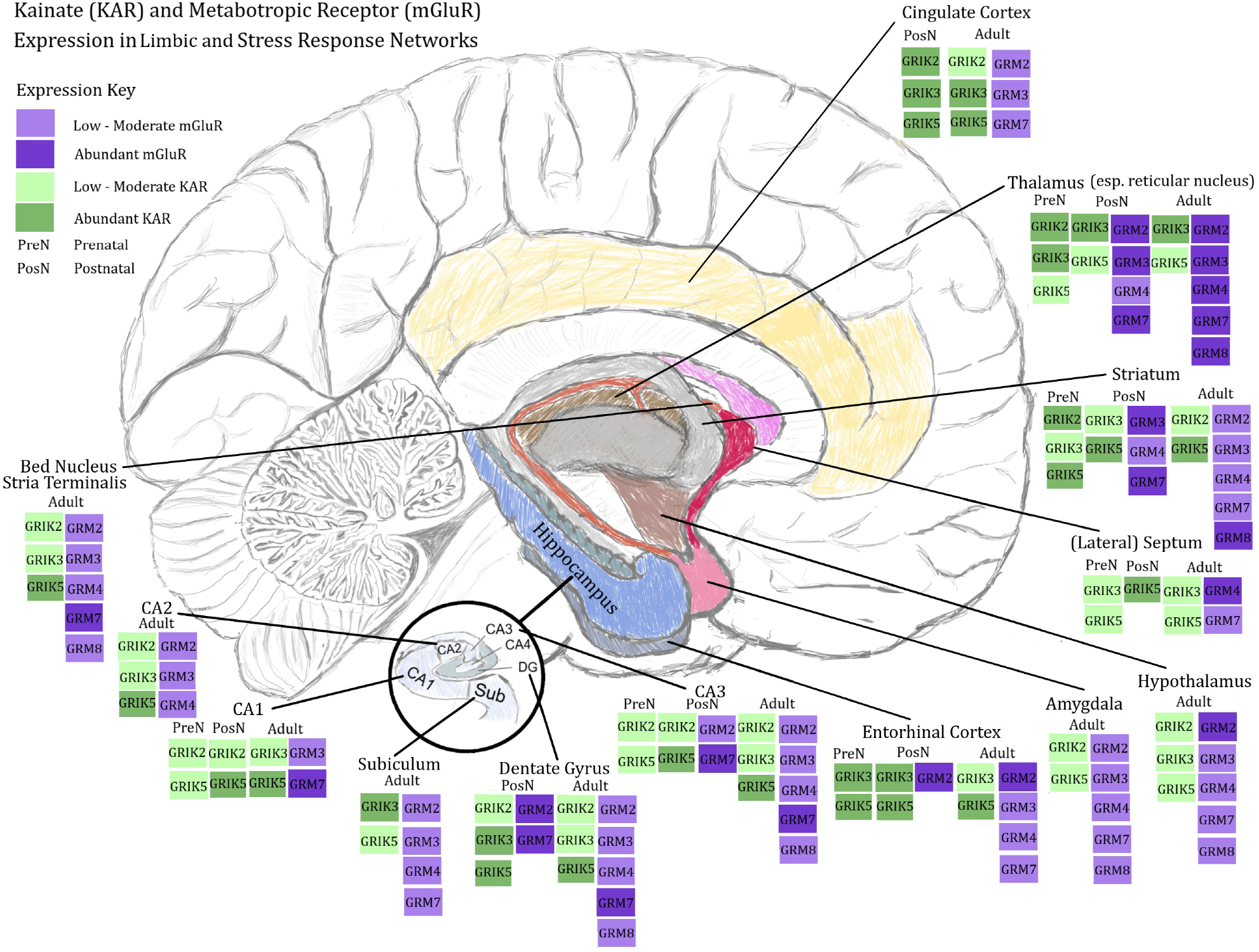
Kainate and metabotropic receptor expression in brain regions crucial for HPA regulation. (Most detailed brain-expression data come from rodent studies. Where available, we have used data from human fetal or postmortem studies. Broadly, there are cross-species parallelisms in kainate and metabotropic expression. Similarities and differences are discussed in Supplementary section S3.)

### Control of HPA function by metabotropic and kainate glutamate receptors

The PVN is the crucial hypothalamic mediator of psychogenic stressors that drive HPA activity. Glutamate acting directly on parvocellular neurons of the PVN stimulates CRH release, whereas GABA inhibits this [7]. This means that modulation of glutamatergic signaling by both kainate and metabotropic glutamate receptors may serve to inhibit direct activation of the PVN.

*GRIK2, GRIK3*, and *GRIK5* are all expressed in the PVN and surrounding regions in adult rats, although *GRIK1* is the most abundantly expressed kainate receptor subunit mRNA in the PVN proper [83]. *GRIK5* is extensively expressed on parvocellular neurons [84]. Presynaptic activation of GRIK1 subunits in the PVN has been shown to modulate HPA activity by inhibiting CRH from parvocellular neurons [62]. Similarly, agonism of presynaptic kainate receptors in hypothalamic neurons facilitates inhibitory GABAergic signaling [85].

*In vitro* antagonism of Group II metabotropic receptors in hypothalamic slices has been shown to increase CRH signaling, whereas no other metabotropic receptor agonists or antagonists had this effect. Mice administered with Group II antagonists *in vivo* experienced an increase in corticosterone that mimicked the response to the forced-swim (behavioral despair) test [86]. Given the predominant presynaptic and glial modulatory functions of Group II receptors, the above evidence implicates Group II receptors in the attenuation of central gluta-matergic inputs to the hypothalamus (likely in the PVN), attenuating the HPA stress response. Group III metabotropic receptors have also been implicated in modulating excitatory inputs to the lateral hypothalamus [87], while Group I metabotropic receptors stimulate both oxytocin and vasopressin release from the SON [88].

Agonism of Group II metabotropic receptors disrupts the fear-potentiated startle response in mice, suggesting that these receptors regulate the learning of fearful experiences [89]. Knockout of *GRM2* has been shown to correlate with increased stress in social interactions [90]. In macaques captured from the wild, six-week chronic intravenous administration of a Group II agonist reduced basal cortisol levels by as much as 50% compared to controls [91]. This same agonist has been shown to act on GRM3 receptors in adrenal gland cells, leading to a reduction in aldosterone and cortisol via inhibition of the adenylyl cyclase / cAMP signaling pathway [92]. Yet another Group II agonist attenuates aggressive tendencies, hyperactivity, and deficits in the inhibition of the startle response of mice reared in isolation [93]

It has been proposed that Group II metabotropic receptors in the central amygdala dampen the stress response by modulating the release of glutamate, in turn leading to an increase in GABAergic signaling and overall dampening of excitatory inputs to the PVN. Agonism of these receptors also leads to an increase in activity in the predominantly inhibitory BNST, and in the the PVN in response to stress, suggesting modulation of the HPA by suppression of excitatory signaling. At the same time, activation of Group II receptors is downregulated in the hippocampus [94]. Excitatory feedback outputs from the hippocampus likely act on inhibitory neurons of the BNST that, in turn, relay to the hypothalamus [82]. Thus, decreased Group II modulation of these outputs in response to stress can have the effect of enabling increases of inhibitory signals to dampen the HPA cascade. Within the BNST itself, activation Group II and III receptors has been shown to suppress excitatory transmission [95].

Mice lacking the Group III receptor *GRM8* display increased age- and sex-dependent anxiety-like behaviors and startle response [96, 97]. However, in contrast to *GRM2*, knockout of *GRM8* can enhance social interactions, suggesting that this receptor has opposing effects depending on the nature of the stressor. In another contrast with *GRM2* and the Group II subfamily as a whole, ablation of *GRM7* can make mice less fearful and less aggressive [98, 99, 100] correlating with a severe reduction in neuronal activity in the BNST. This suggests that GRM7 serves to enhance overall excitability of BNST neurons projecting to the PVN (and thus the stress response), perhaps through modulation of glutamatergic release innervating GABA inhibitory interneurons [100]. Activation of GRM4 reduces anxiety-like behaviors in mice, while knockout enhances fear-conditioning responses and increases anxiety in adult but not juvenile mice [101]. Such anxiety-related effects are thought to be brought about by alterations to amygdalar function, whereby either excitatory or inhibitory signaling is modulated by Group III receptors.

Although the above behavioral correlates of Group II and III receptor (ant-) agonism and ablation are partially contrasting, they indicate that both subfamilies are important for modulating the stress response, including aggressive reactivity. This tendency is clear in Group II metabotropic receptors, while the actions of Group III receptors are more varied according to the specific subunits and brain regions activated. Studies in rodents suggest that GRM8 and GRM7 have broadly opposite effects on anxiety levels, with GRM8 activation tending to be closer to Group II metabotropic receptors in its anxiolytic effects, while GRM7 seems to be more anxiety- (and aggression-) inducing [102]. This suggests that the numerous signals of selection on *GRM8* across domesticates and similar signals on Group II receptors in humans may be markers of convergent selection for a decreased stress response. This said, evidence also points more clearly towards activation of Group II receptors in potentiating prosocial behaviors. Future investigation of differences in brain region expression of Group II and III receptors in domesticated species may help to shed more light on their contributions of each to the modulation of the stress response.

The kainate receptors we have examined here are abundantly expressed in limbic regions that modulate HPA-axis function via glucocorticoid (GC) feedback (in particular the hippocampus and medial prefrontal cortex, but also more moderately in the amygdala [see Figure 2]). GCs promote glutamate release in these feedback regions, and the different affinities of mineralocorticoid receptors (MRs; bound by GCs at low concentrations) and glucocorticoid receptors (GRs; bound at higher concentrations) enable the modulation of stress feedback signaling from basal or moderate to acute levels [67, 68].

Glucocorticoids differentially modulate the expression of kainate receptor mRNA in the hippocampus depending on whether MRs or GRs are bound [103, 53]. Adrenalectomy (lowering corticosteroid levels) leads to increased expression of *GRIK2* in DG and CA3, and of *GRIK3* in DG [53] (although no change has also been reported for *GRIK2* in CA3 [103]). Single dose treatment with low levels of corticosterone following adrenalectomy — thought to bind MRs — increases *GRIK3* and high affinity subunit (*GRIK4* and *GRIK5*) mRNA in DG, as well as *GRIK5* across the hippocampus [103]. MR binding has been reported both to lower and raise *GRIK2* levels in the hippocampus [53, 103].

Acute corticosterone treatment in adrenalectomized rats lowers kainate receptor mRNA expression to levels of untreated controls [103]. Similarly, chronic treatment leads to lower expression of *GRIK3* and *GRIK4* in hippocampal structures, although no changes were noted for *GRIK2* or *GRIK5*[53]. These divergent MR/GR-mediated patterns of expression can help to elucidate the mechanism by which the genes under selection in domesticates and humans are expressed in a manner that can modulate the stress-induced feedback response.

Kainate receptor activation at CA3-CA1 synapses serves to inhibit glutamate transmission via G_i/o_ signaling, especially when synapses are immature. At mossy fiber synapses connecting DG and CA3 (areas of high kainate receptor expression throughout life), kainate receptors inhibit glutamatergic signaling when glutamate is released at high levels, while facilitating release at lower levels, again via a G_i/o_-coupled mechanism [104]. Similar biphasic modulation has been detected in the neocortex and amygdalae of rodents. Thus, when GCs are circulating at low levels, during basal or low stress, MR binding should lead to higher kainate receptor expression and facilitation of glutamatergic signaling in feedback regions. At higher GC levels (as when under acute stress) GR binding will tend to reduce kainate receptor expression, thus diminishing these receptors’ potency in modulating glutamate release. In contrast, postsynaptic expression of AMPA and NMDA is enhanced under acute stress and corticosterone treatment [105].

Because there are no direct hippocampal, prefrontal, or amygdalar connections to the PVN, feedback from these regions are instead relayed via the BNST, lateral septum, and ventromedial hypothalamus (VMH), which are all predominantly GABAergic [106, 7]. For the amygdala, which emits primarily inhibitory outputs to intermediary regions, this results in “GABA-GABA disinhibitory” downstream signals, increasing excitatory inputs to the PVN [69]. In the case of kainate receptor expression, which is predominant in the hippocampus and medial prefrontal cortex, increased facilitation of glutamate release during basal or low-level stress is likely to primarily innervate GABAergic neurons along pathways relaying to the PVN. Similarly, downregulation of kainate receptor expression by GR binding during acute stress serves to diminish the alternate modulatory effect of kainate receptors during intense glutamatergic release. This, in turn, should allow for NMDA and AMPA receptor signaling to be potentiated, leading to a dampening of the HPA stress response, again via the innervation of inhibitory neurons of the BNST, septum and VMH, which relay to the PVN.

In contrast to prefrontal and hippocampal feedback regions, the effect of glucocorticoids on parvocellular and magnocellular neurons of the hypothalamus is to downregulate glutamatergic signaling via the release of endocannabinoids, in turn promoting the release of GABA [107]. Kainate receptors have been implicated in the mobilization of endocannabinoid signaling in distinct brain regions, as well as in the promotion of GABAergic signaling in the PVN [62, 108, 109].

Figure 3 presents a schema of the modulatory actions of metabotropic and kainate receptors in the stress-response cascade.

**Figure 3:**
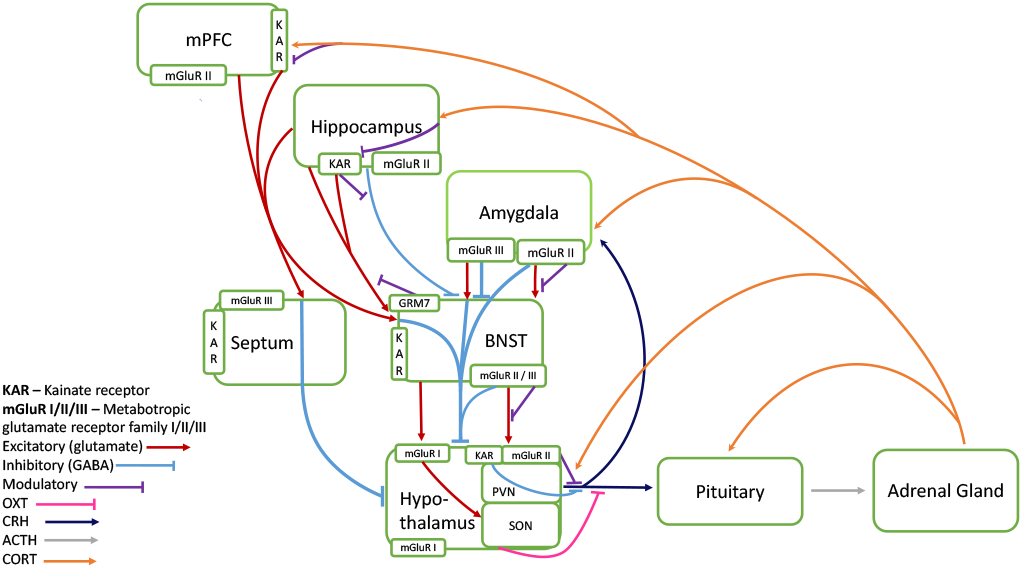
Modulatory Actions of Metabotropic and Kainate Receptors

Increased metabotropic and kainate-mediated attenuation of central and feedback stress responses may plausibly have conferred selective advantages in human evolution, not only via the reduction of stress and enabling of prosocial cooperation, but also by enabling subsequent increases in *GRIK2, GRIK3*, and *GRIK5* expression: Firstly, Group II and III metabotropic glutamate receptor modulation of amygdalar fear processing in response to stressors, in combination with kainate and Group II metabotropic receptor inhibition of CRH release in the PVN can lead to a signaling cascade that results in lower glucocorticoid feedback in limbic structures. This, in turn may lead to increased expression of kainate receptors in the hippocampus, prefrontal cortex, and elsewhere, enabling subsequent selection on improvements in plasticity and learning, as well as further resources for limbic modulation of the stress response.

## 4 Conclusion

We have argued here that, on balance, selective pressures have led to a modulation of glutamatergic signaling in order to attenuate the HPA stress response, and that this has had the corollary effect of increasing synaptic plasticity in limbic feedback regions that play a crucial role in memory and learning. We have not dealt in detail with G-protein signaling cascades activated by kainate, Group II, and Group III receptors. A more in-depth view of convergence may be gained by exploring the extent to which the same signaling cascades (in particular G;/o signaling) are activated in homologous brain regions across species regardless of the receptor subtype initially bound.

Given the important roles that serotonin and oxytocin play in promoting social and empathetic behaviors across different species, it has been proposed that convergent tameness in domesticates and prosociality in modern humans is driven by alterations to these systems [3]. The present analysis of modern human-archaic and domesticate-wild differences in glutamatergic receptor genes suggests that any such modifications (particularly to oxytocinergic signaling) are more likely to be dependent on upstream changes to glutamatergic signaling. Glutamate mediates the release of oxytocin and vasopressin from the SON and PVN [110].

Serotonin modulates glutamatergic activity in the brain, often inhibiting excitatory potentials and stimulating GABAergic inhibitory signaling [111]. This regulation from outside the glutamatergic system could potentially produce comparable modulation of the stress-response to the regulation from within that we propose for metabotropic and kainate receptors. In studies of genetic differences between tame and aggressive foxes, changes to serotonergic signaling accompany those of the glutamatergic system, with genetic changes often potentially relevant to both synapses [30, 31]. Similarly, domestication of the pig appears to have involved important changes across both systems [39]. Signals of selection associated with serotonergic signaling have also been identified in a tame strain of rat and the domesticated goat [112, 35]. It should be noted, however, that glutamatergic signaling can also control the release of serotonin from the dorsal raphe nucleus [113].

One may reasonably ask whether Group II and III metabotropic and kainate receptors share functional or structural qualities that have made them more likely to come under selection than NMDA, AMPA, or Group I metabotropic receptor (sub)families in domesticate species and modern humans. These receptors, particularly NMDA, have been implicated in many disorders and phenotypes reviewed here for metabotropic and kainate receptors [114, 115, 116, 117, 118, 119, 120, 63]. Even within receptor families, specific subunits (and combinations of them) or splice variants can have diametrically opposing functional properties from others that are nominally similar [121, 122].

The similar modulatory properties, overlapping expression patterns, and shared phenotypical associations of those genes that are most consistently detected across domestication and modern human studies has led us to hypothesize an overarching role for kainate and Group II and III metabotropic receptors in the modulation of the stress response. However, there is no reason, in principle, why other glutamate receptor families could not have contributed to the emergence of tame behaviors. In fact, signals are detected on NMDA receptors in two species for each of *GRIN2B* (fox and yak), *GRIN2D* (AMH and fox), and on three species in the case of *GRIN3A* (AMH, yak, and duck; see Table 2). Each of these genes’ respective subunits are highest expressed during embryonic and early postnatal stages, and have been associated with the maintenance of immature dendritic spines during development [123] If similar patterns of selection should be detected on NMDA receptors across other domesticates, a case could be feasibly be made for their contributing to the juvenile cognitive phenotype typically retained by domesticate species into adulthood. On the other hand, the only case of potentially convergent selection on an AMPA receptor gene occurs on *GRIA2* (AMH, cat, and pig), which is a crucial AMPA subunit at mature synapses [124].

The diversity of roles played by kainate receptors in the CNS, including extensive excitatory, inhibitory, ionotropic, metabotropic, and modulatory activities [65], provides a wide range of possible functions upon which natural selection can act. Selective pressures seem to have honed varied specializations for kainate receptors in different brain regions. Group II and III metabotropic receptor subunits can have opposing actions within a single brain region, and to stressful stimuli being processed. This suggests that these subunits often have complementary modulatory functions within their subfamilies. Nonetheless, there are considerable overlaps of function, and these subunits primarily inhibit adenylyl-cyclase signaling. Each receptor subfamily acts, broadly, to dampen stress responses. These overlaps in function undoubtedly contribute to numerous signals of selection being spread across different metabotropic receptor genes in distinct species.

In the future, it may be worthwhile examining the extent to which the glutamatergic changes discussed here could have impacted the vocal abilities of the relevant species, including ours. Although the extent of archaic humans’ vocal-learning abilities is not known, this capacity is highly striatum-dependent, as can be seen in the neurology of Tourette’s syndrome. Glutamatergic signaling alterations in the striatum have been implicated in Tourette’s syndrome, including genes with AMH-specific changes. Vocal learning deficits in humans who carry a mutation on the *FOXP2* gene are thought to arise, in part, from abnormalities in the striatum [125]. Knock-in experiments of the humanized *FOXP2* allele in mice have shown the principal structural changes to take place in the striatum, where MSN dendrites are longer and respond to stimulation with increased long term depression [126]. *FOXP2* is also expressed in glutamatergic projection neurons of the motor cortex and is highly expressed during the development of corticostriatal and olivocerebellar circuits, important for motor control [126, 127]. Several glutamate receptor genes discussed above, such as *GRIK2* and *GRM8*, have been identified as a transcriptional target of FOXP2 in the developing human brain [128]. Significant changes to glutamatergic expression have also been found in the vocal nuclei of vocal-learning birds, and in the domesticated bengalese finch compared to its its wild ancestor the white-rumped munia [9, 129, 130]. These changes are thought to play a role in the more complex singing capabilities of the bengalese finch. Glutamate receptor genes form part of the most highly co-expressed module in the periaqueductal gray of bats, a mammalian species that displays strong evidence of vocal-learning capabilities [131].

Finally, an important question arising from the evidence presented above is how glutamatergic involvement in the modulation of the stress response relates to the hypofunction of the neural crest, proposed to account for the suite of phenotypic changes that make up the domestication syndrome [17]. At this point, the evidence from selective sweep studies strongly points to the involvement of glutamatergic signaling in domestication. Our comparison of different receptor types does not allow evaluation of arguments that a “mild neurocristopathy” drives the emergence of the domesticated phenotype. One neural crest-related gene (*KIT*) was detected in studies of cattle, sheep, pigs, and yaks. This gene is involved in melanocyte differentiation, and evidence points to its importance in selection for coat colour: Signals occurring on *KIT* in Clydesdale horses and Birman cats have been implicated in their breed-specific coat patters [29, 28]. As such, selection on *KIT* is unlikely to be related to the unifying domesticated trait of tameness, as proposed here for modulatory glutamatergic signaling. Whether this extends to other neural-crest related genes, in line with arguments against the universality of the effects predicted by the neural crest hypothesis (see [132]) remains to be determined.

There are various possible explanations for how glutamatergic signaling may interact with, or even bypass, the neural crest to bring about the broader phenotypical features of the domestication syndrome. We present two alternatives in Supplementary Section S4 that future work could put to the test. On considering these two alternative accounts of glutamatergic signaling in the emergence of the domestication syndrome (NCC-interacting versus NCC-independent), we prefer to be cautious about attributing too many functional consequences to the signals of selection highlighted here. We consider that a mild neurocristopathy will almost certainly explain some physical trait changes in domesticate species, and that this may serve to entrench earlier selection for tameness via reduced inputs to stress-hormone cells in the adrenal glands. Future investigations may help to determine whether these reductions result from upstream modulation of glutamatergic signaling. In some species, changes to glutamatergic signaling may also have acted to directly alter the development of physical traits, independently of genetic or epigenetic alterations to the neural crest, driven by domestication.

## 5 Materials and Methods

Based on our previous work showing overlapping signals of selection on glutamatergic-signaling genes in domesticates and anatomically modern humans [4], we extended our comparative analysis across a broad range of domesticated species. In order to delimit the comparison to a clearly-defined gene set within the glutamatergic system, we selected the 26 glutamate receptor genes as our cross-species comparator (Table 1).

For our comparison, we included all domesticated species for which whole-genome sequences were available. In species for which no studies of selective sweeps were available, we included studies of decreased heterozygosity differentiating tame and aggressive strains (fox) and/or studies detailing brain-expression differences between tame and wild or aggressive lineages (guinea pig, fox, and rat). We also included a study of introgressed genes from cattle to yaks, which identified genes likely to have been subject to positive selection. Finally, we included for comparison high-frequency (near-fixed) changes differentiating modern from archaic humans. (Table 2.)

We extracted all instances of selective sweeps, introgression, brain-expression, and high-frequency changes on glutamate receptor genes from the domestication and modern-human studies listed in Table 2. We compiled these genes for comparison according to the receptor (sub)families of which they are members. To evaluate whether signals detected on glutamate receptor genes occurred at a rate significantly higher than those for comparable receptor classes, we compiled signals of selection occurring on ligand-gated ion channel receptor and G-protein coupled receptor genes. We excluded olfactory, vomeronasal, and taste receptor genes from our comparison, given the large cross-species variability in functioning receptors encoded by these genes, driven, in large part, by species-specific dietary and environmental specializations. We also excluded orphan g-protein coupled receptors from our analysis, given that, for the most part, their endogenous ligands and broad functions remain uncharacterized. Because of an earlier observation that domestication events may have affected Erb-b2 signaling [4], as well as extensive evidence for effects on stress-hormone levels, we also included receptor kinase and nuclear hormone receptor genes for comparison.

Following the compilation of signals on 488 receptor genes across thirty domestication studies, we confirmed that this data, when categorized according to receptor subfamilies, passed the assumptions required for use of a one-way ANOVA (residuals-versus-fit and Levene’s test for homogeneity of variance). We then carried out the one-way ANOVA in R, showing significant differences in signals of selection between certain receptor subfamilies. Following this, we used Monte Carlo random sampling (100,000 simulations) to determine the chance likelihood of signals occurring at the rates they do on single genes and gene (sub)families.

In order to determine potentially shared functions of kainate and metabotropic receptor gene (sub)families that showed the strongest signals of convergence across humans and domesticates, we investigated associations with phenotypes relevant to tameness and prosociality by doing an exhaustive literature search (Pubmed). These included associations with the startle response and neurodevelopmental, neuropsychiatric, stress, and mood disorders, summarized in Table 3.

Finally, we carried out a meta-analysis (summarized in Figure 2, details in Supplementary Material) to determine the brain expression of kainate and Group II and III metabotropic receptor genes.

## Author contributions

Data collection and analysis: TOR, CB; proposed mechanism: TOR; draft preparation: TOR, with input and revisions from CB; project supervision: CB.

## Acknowledgements

We would like to thank Pedro Tiago Martins for invaluable help in preparing Tables 1 and 2, and Figure 1. We kindly thank Bridget D. Samuels and Kay Sušelj for help in choosing and carrying out the statistical analyses. We thank Yan Li for sharing unpublished data relevant to the study in [23]. We would also like to thank Adam S. Wilkins, Stefanie Sturm, Alejandro Andirkó, and Pedro Tiago Martins for helpful comments on earlier drafts of this text.

