## Supplementary Material for "Converging roles of glutamate receptors in domestication and prosociality"

### Supplementary Material: Converging roles of glutamate receptors in domestication and prosociality

[1]Universitat de Barcelona
[2]Universitat de Barcelona Institute of Complex Systems
[3]ICREA

February 1, 2019

#### S1 Broader changes to glutamatergic signaling under domestication

Signals of selection have been detected in AMH and Yaks on *NETO2* [1, 2], which encodes an accessory subunit that modulates the desensitization of kainate receptors, and interacts with the GRIP scaffolding protein to increase receptor abundance at synapses [3, 4]. In mouse hippocampal cultures, NETO2 and GRIK2 proteins interact to increase recycling of the cotransporter SLC12A5 to the surface membrane [5] (signals of selection on the corresponding gene in AMH [6]). Increased abundance of SLC12A5 at hippocampal dendritic spines suppresses AMPA mediated excitatory transmission by preventing the transmembrane diffusion of GRIA1 proteins [7], and NETO2-SLC12A5 interactions during development promote increases in GABAergic inhibitory signaling in the hippocampus [8]. Altered prefrontal expression of different *SLC12A5* transcripts have been associated with schizophrenia and affective disorders [9], and this gene has also been associated with autism, epilepsy and developmental delay [10, 11, 12]. The signals of selection on *NETO2* and *SLC12A5*, their postsynaptic interactions with kainate receptors, regulation of excitatory and inhibitory transmission, and associations with affective and developmental disorders make them candidate interactors for the modulation of the stress response in both domesticate species and humans.

In a recent study of convergent signals of selection on domesticated goats and sheep, Neurobeachin (*NBEA*), a gene that regulates glutamate and GABA receptor expression at synapses, was highlighted as being one of the most likely genes to be implicated in behavioral changes under domestication [13].

In the case of rats selected for tame behaviors, expression differences are also prominent in glutamatergic-signaling genes implicated in the stress response. Most notably, *SLC17A7*, which encodes for a vesicular glutamate transporter, is expressed twice as abundantly in the brains of tame rats as in those of the aggressive strain. Underexpression of *SLC17A7* has been implicated in increased anxiety in mice [14].

The postsynaptic scaffolding proteins DLGAP1 and DLGAP3, have both been implicated in OCD and Tourette's Syndrome [6, 1, 15, 16, 17]. *DLGAP1* and *GRIK2* have been identified in a genome-wide association study as high-confidence interactors in the etiology of OCD [18]. Signals of selection have been identified on *DLGAP1* in a horse domestication study [19]. Stereotyped and compulsive behaviors are common in horses, as well as many other domesticated animals [20, 21]. Indeed, pharmacological targeting of glutamatergic signaling is successful in treating both Canine Compulsive Disorder and human OCD, suggesting a shared etiology for these conditions [22]. Both *DLGAP1* and *DLGAP3* (signals of selection in AMH [1, 6]) have been implicated in Tourette's syndrome and OCD, as well as schizophrenia and major depression. At the postsynaptic density, DLGAP proteins interact with DLG membrane-associated proteins, and associate with kainate receptors. Signals of selection have been detected in on *DLG4* in yaks [2]. When cotransfected in cell cultures, GRIK2 subunits associate with DLGAP3 only when accompanied by DLG4 [23]. GRIK2/GRIK5 heteromeric receptors desensitize more slowly to glutamate when they associate with DLG4 [24].

*DLGAP1* knockout mice exhibit marked social deficits, correlating with loss of associations between postsynaptic scaffolding complexes and the DLG membrane-associated proteins [25]. *DLGAP3* knockout mice exhibit OCD like behaviors, and have increased NMDA and reduced AMPA transmission at corticostriatal synapses [26]. Knockout of all five kainate receptor genes (*GRIK1-5*) results in comparable preservative behaviors, yet also leads to reduced NMDA: AMPA ratios at corticostriatal synapses [23]. The same effects were not found in CA3 of the hippocampus, suggesting that kainate receptors are not essential for NMDA mobilization in this region. Postsynaptically localized GRIK2 receptors inhibit glutamate signaling in striatal direct pathway medium spiny neurons by activating presynaptic cannabinoid $CB_1$ (G-protein-coupled) receptors, thus regulating plasticity in basal ganglia circuits [27]. The interactions of kainate receptors with DLG and DLGAP proteins, their roles in the regulation of striatal plasticity, and their involvement overlapping stress-related phenotypes make them plausible candidates for having come under convergent selection in domestication and modern human evolution.

Many glutamate signaling genes have been identified in a recent study of genomic regions associated with tame behaviors in the domesticated fox [28]. These include *SORCS1*, whose encoded protein is involved in trafficking and maintaining AMPA receptors at the cell membrane, increasing synaptic transmission [29]. This gene was detected alongside other glutamatergic signaling genes, including *SORCS2*, *SORCS3* (identified in the fox study), and the AMPA receptor gene *GRIA1* in a draft human selective sweep study [30]. However, in a revised version of the same paper, only signals on *SORCS2* remained [6]. The SORCS2 protein is highly enriched on post-synaptic densities in the striatum and hippocampus, where it regulates NMDA-dependent synaptic plasticity and spatial learning [31]. *SORCS2* knockout mice display manic-depressive symptoms, and polymorphisms in the human population have been associated with bipolar disorder, schizophrenia, and attention deficit hyperactivity disorder (ADHD) [32]. *SORCS3* knockout mice have fear-extinction and spatial-memory deficits [33]. Each of the above SORCS proteins interact with DLG4 [29, 31, 33].

*PLCB1* and *PLCB4* display signals of selection in red foxes, pigs, and yaks [28, 34, 2]. PLCB4 acts downstream of GRM1 synaptic signaling to mobilize endocannabinoids, which suppress glutamate release [35]. In humans, this gene has been implicated in syndromic malformations of the ear and jaw [36]. PLCB1 also modulates GRM signaling and has been implicated in schizophrenia [37, 38]. Signals on *KCNJ3* have been identified in foxes and multiple human selective sweep studies [1, 30, 15, 28]. This gene is expressed in the adrenal gland, and its encoded protein has been implicated in cell guidance and localization during CNS development, associates with postsynaptic glutamatergic proteins, and has been implicated in schizophrenia [39, 40, 41].

*CACNA1C*, which encodes for a voltage-dependent calcium channel subunit, has been detected in a cattle domestication study, and distinct allelic variants are associated with tame and aggressive breeds of foxes [42, 28]. In humans, this gene has been associated with bipolar disorder, schizophrenia, and major depression [43]. A single polymorphism on *CACNA1C* in schizophrenics is associated with increased expression and higher activity in the hippocampus in response to emotional stimuli [44], as well as higher left amygdala activity in bipolar disorder [45]. *CACNA1D* has been detected in human and horse selective sweep studies [1, 19]. Both *CACNA1C* and *CACNA1D* are abundantly expressed in striatal medium spiny neurons (MSNs) [46]. CACNA1D subunits bind to glutamatergic scaffolding proteins of striatopallidal MSNs, where they inhibit glutamate transmission by allowing $Ca^{2+}$ influx, thus downregulating the dendritic spine densities. MSNs are targeted by dopaminergic afferents which drive glutamatergic transmission and inhibit CACNA1D channel opening. When dopaminergic signaling is abolished, spine densities are decreased via the opening of CACNA1D channels in a process thought to be critical in the motor deficits experienced by Parkinson's disease patients [47]. *CACNA1D* is also expressed in the adrenal gland [48].

The abundant striatal expression of kainate and other glutamatergic signaling genes, their interactions at synapses, overlapping involvement in cognitive and stress disorders, including OCD, and associations with tame behaviors in the fox, suggest that convergent selection on striatal-mediated behaviors may have accompanied modulation of the HPA stress response. In the case of humans, it has recently been argued that increases in prosociality following our divergence from other extant primate lineages was driven by neurochemical changes, especially to dopaminergic signaling, in the striatum [49]. We propose that such selective pressures on prosociality have continued following our split from Neanderthals, and that convergent signals of selection with domesticates across the glutamatergic system are indicative of this.

*SLITRK1*, which encodes for a transmembrane protein at excitatory synapses, has a high-frequency missense change in modern humans and may be under positive selection in our lineage [50, 6, 51]. This gene has also been implicated in OCD and Tourette's syndrome [52]. Beyond the receptor genes highlighted in the present study, OCD is associated with a number of genes related to glutamatergic signaling that show signals of selection in AMH. These include *SLC1A1*, which encodes for a neuronal glutamate transporter, is expressed prominently on hippocampal dendrites, and is most consistently associated with OCD among glutamatergic signaling genes [6, 53, 54, 55].

### S2 Kainate and metabotropic receptor genes in human and domesticated behavioral phenotypes

#### Kainate receptor genes in human and domesticated behavioral phenotypes

*GRIK3* has been implicated in artificially selected behavioral differences between modern domesticated cattle breeds: Lidia cattle (Spanish Fighting Bulls), which have been artificially selected for agonistic behaviors, display differential signals of selection on *GRIK3* when compared to tame breeds [56]. These allelic differences, in turn, implicate ancient signals of selection detected on *GRIK3* in the emergence of the domestic phenotype in cattle [42], resulting from selection for tameness.

Similar functional implications are likely for yaks, for whom signals on *GRIK3* have been detected from cattle introgression. *GRIK3* has also been identified in various dog domestication studies (see Table 2). As in cattle, *GRIK3* is a strong candidate for having been subject to changes in the initial stages of domestication [57].

In humans, *GRIK3* deletion has been implicated in severe developmental delay affecting language, motor, social, and attentional skills [58], as well as a a range of neuropsychiatric disorders and related phenotypical traits. These include schizophrenia [59], bipolar disorder (manic-depressive illness) [60], major depression [61] (including significantly higher expression levels in the dorsolateral prefrontal cortex in cases of suicide [62]), high anxiety comorbid with depression [63], increased startle response [64], and personality traits such as decreased cooperativeness and compassion, and increased shyness and harm avoidance [65].

*GRIK2* has been implicated in many of the same neuropsychiatric and neurodevelopmental disorders as *GRIK3*, including schizophrenia [66, 67], bipolar disorder [68], autism spectrum disorder [69], and major depression (including emergent suicidal ideation during treatment for the disorder) [70, 62]. Both genes code for low-affinity kainate receptor subunits which share many brain-region and synaptic localizations, and which can coassemble with each other or with high-affinity kainate receptor subunits [71, 72, 73]. These similarities likely explain why *GRIK2* and *GRIK3* are implicated in many of the same clinical disorders, and suggests that changes to these genes may have played similar roles in domestication and recent human evolution.

Deletions and loss-of-function mutations on *GRIK2* can cause intellectual disability and social impairments, including autistic behaviors [74, 75, 76]. A gain-of-function mutation on *GRIK2* in humans has been shown to bring about syndromic changes during development that include slight microcephaly, intellectual disability, happy demeanor, short attention span, increased drooling, stereotyped behavior, and increased aversion to loud noises [77]. Some of these symptoms are comparable to behaviors and phenotypes often observable in domestic dogs.

The Czechoslovakian Wolfdog, a tame hybrid resulting from a recent wolf-dog crossing experiment, has accumulated a significantly higher number of dog-like polymorphisms on *GRIK2* [78], making this gene a strong candidate gene in enabling the retention of tame behaviors. Domestic rabbits — which have signals of selection on *GRIK2* — exhibit gross brain changes in regions known to mediate fear response [79].

*GRIK2* knockout mice experience behaviors reminiscent of manic episodes in humans, including decreased manifestations of fear, anxiety and despair in response to stressful situations, as well as increased risk-taking and aggressiveness towards conspecifics [80, 68]. Among humans, aggression and antisocial personality disorder are significantly higher in bipolar and schizophrenic patients than in the wider population [81, 82, 83, 84].

Polymorphisms on *GRIK2* have been associated with obsessive compulsive disorder (OCD), a stress disorder, but there is no strong evidence for the involvement of *GRIK3* as yet [85, 86]. Mice that have all five kainate receptors knocked out exhibit increased preservative and diminished exploratory behaviors, a tendency typical in OCD [23].

Significantly lower expression of *GRIK5* has been reported in the hippocampus, prefrontal cortex, and various thalamic nuclei of schizophrenia patients, [66, 87, 88]. A single nucleotide polymorphism (SNP) on this gene has been implicated in bipolar and panic disorders, and rare missense mutations have been associated with autism spectrum disorder [89, 90]

#### Metabotropic receptor genes in human and domesticated phenotypes

There is ample evidence that metabotropic glutamate receptors are implicated in the emergence of tame and prosocial behaviors, in modulating the stress response, and in many of the same neurodevelopmental and neuropsychiatric disorders as kainate receptors.

A fixed missense mutation on *GRM3* differentiates aggressive farm-bred foxes from their tame counterparts and is thought to be one of the most important signals of the behavioral differences between the two groups [91]. There is experimental evidence that GRM3 activation in the medial prefrontal cortex is necessary for fear extinction via long-term depression of synaptic connections [92], and agonism of Group II metabotropic receptors inhibits the expression of fear potentiated startle response in rats [93]. Group II metabotropic receptors are implicated in stress and mood disorders including major depression, anxiety disorders, and obsessive compulsive disorder, as well as schizophrenia and bipolar disorder [94, 95, 96, 97, 98, 99]. *GRM3* has been identified as having longer homozygous segments in autism spectrum disorder (ASD) subjects than their parents [100], and activation of Group II receptors can help to ameliorate ASD-like social deficits in mice [101]. Agonism of Group II receptors also serves to attenuate attention deficits and impulsivity brought about by overactivation of the 5HT2A serotonin receptor in a rodent model of ADHD [102].

The Group III metabotropic receptor gene *GRM8* is detected in more domesticate or human selective sweep studies than any other metabotropic receptor gene. In one natural experiment, very much relevant to domestication but not encompassed by these selective sweep studies, the endangered and highly inbred Apennine brown bear, known for its docility towards humans, has significantly lower nucleotide variability on *GRM7* and *GRM8* than the genome-wide average. Similar patterns were detected on other genes related to tameness in domesticated species. The authors of the study postulated that these changes were the result of selection on reduced aggressiveness in the Apennine bear population [103].

In humans, polymorphisms on Group III receptors are often associated with mood and stress disorders, including anxiety and major depression [94, 104, 105, 106]. Studies in rodents broadly seem to support the view that activation of GRM8 serves primarily to reduce anxiety, whereas GRM7 activation has the opposite effect [94]. *GRM8* knockout mice have generally increased levels of anxiety, although some measures are age- and sex-dependent, and social anxiety is attenuated by knockout [107, 108]. On the other hand, *GRM7* knockout broadly decreases anxiety in mice [109].

Knockout of *GRM7* causes fear-response and fear-memory deficits, while agonism of GRM8 receptors in the amygdala inhibits acquisition and expression of fear as measured by startle response [110, 111]. Polymorphisms or partial duplications on *GRM7* and *GRM8* have been associated with schizophrenia, ADHD, major depressive disorder, and autism spectrum disorder [112, 113, 114, 115, 116, 104, 117, 118]. *GRM7* alone has been associated with bipolar disorder, while *GRM8* has been implicated in severe intellectual disability [119, 120].

Polymorphisms on *GRM4* have been associated with bipolar disorder, schizophrenia, and major depressive disorder [121, 106]. Activation of this receptor subunit can relieve autism-like symptoms and reduce anxiety-like behaviors in mice, while knockout enhances fear-conditioning responses and increases anxiety in adult mice [122, 123]. Potentiation of GRM4 receptors reduces certain measures of impulsivity, while potentiating others and impairing visual attention (ADHD-like phenotype) in rats [124].

ADHD is often comorbid with obsessive compulsive behaviors and Tourette's syndrome, both of which have been linked to aberrant glutamatergic signaling, the former being associated with both kainate and Group II metabotropic receptors [125, 126]. Group III metabotropic receptors have been proposed as potential targets for treatment of OCD and Tourette's syndrome [127].

### S3 Glutamate receptors in the CNS

#### Kainate receptor expression

Although kainate receptors are structurally similar to AMPA receptors, "they fulfill a more varied range of synaptic and extrasynaptic roles" than their ionotropic counterparts [27]. Apart from ligand gating, kainate receptors modulate glutamate release and activate second-messenger signaling cascades to a greater extent than AMPA or NMDA receptors [128, 129, 130]. Kainate receptors may serve to facilitate or inhibit glutamate or GABA release, regulate dendritic outgrowth during development, potentiate AMPA-dependent excitatory synaptic transmission, or activate distinct "non-canonical" (non-ionotropic) signaling, including $G_{i/o}$ proteins, the principal second-messenger signaling targets of Group II and III metabotropic receptors [131, 132, 133, 130]. Thus, depending on the primary roles played by kainate receptor subunits in a given brain region, their expression may have distinct functional implications far beyond excitatory synaptic transmission [130].

Here, we focus on the regional expression of the low affinity receptor genes, *GRIK2* (because of signals of potential convergence among domesticates) and *GRIK3* (because of parallel signals among domesticates and humans), as well as the high-affinity receptor gene *GRIK5* (given signals of selection detected in AMH). Because signals of convergence with domesticates have yet to be identified for *GRIK5*, this receptor gene received little attention in the cross-species comparisons of selective sweep studies. However, *GRIK5* has been identified in both selective-sweep studies of modern human populations and among high-frequency differences distinguishing archaic from modern human populations [6, 51]. For these reasons we include it in our review of expression data.

*GRIK2* mRNA is highly expressed in cerebellar granule cells and is also found in the caudate putamen and hippocampus, while *GRIK3* is strongest expressed in deep layers of the cerebral cortex, the cingulate cortex, the subiculum, the caudate-putamen, thalamic reticular nucleus, and stellate/basket cells of the cerebellum [134, 73]. *GRIK5* is the most universally expressed of all kainate receptor genes and can be detected across most brain regions, although the encoded subunit must coassemble with low-affinity kainate receptor subunits (GRIK1-GRIK3) in order to form functional ligand-gated ion channels [135, 134, 73]. In general, kainate receptor gene expression is highest in the hippocampus and the cerebellum [73]. Kainate receptor mRNA is detectable in the amygdala, although *GRIK1* is considerably more abundant than other receptor genes that have been detected, with *GRIK2* having moderate and *GRIK5* low expression [136].

Expression levels of kainate receptor genes are developmentally regulated, peaking in the rat brain in the late embryonic and early postnatal period [134]. As early as embryonic day 10, four kainate receptor genes (*GRIK2*-*GRIK5*) are expressed in the rat neural tube. This expression is higher than that of AMPA receptors and is present in both progenitor cells and differentiated neurons, with GRIK2 and GRIK5 subunits being upregulated during neural cell differentiation [137]. During later embryonic periods, brain expression is highest in the cerebellum and striatum, later extending to the cortex, septum, hippocampus, and thalamus. Expression levels drop markedly in the thalamus from the late postnatal period onwards [134]. *GRIK5* is highest expressed in the cortical plate in the late embryonic period. In general, *GRIK2* is not highly expressed in the cortex, although it is abundant in the cingulate cortex, peaking during early postnatal development. *GRIK3* is more broadly expressed in the surrounding cortex than *GRIK2*, but autoradiograph images nonetheless show increased expression in the cingulate cortex during the immediate postnatal period, and moderate to abundant expression into adulthood [135, 134]. Highest expression levels for *GRIK3* during postnatal development are detected in the entorhinal cortex [134].

In the rat hippocampus, expression of kainate receptor genes peaks in the late embryonic and early postnatal period. This is particularly transient in CA1 but remains high in CA3 into adulthood. Expression of kainate receptor genes in the dentate gyrus (DG) also remains high from the postnatal period into adulthood. Similarly, immunolabeling of GRIK2, GRIK3, and GRIK5 subunits in adult rats show highest abundance in the pyramidal layer of CA3 and in neurons in DG, being present in both cell bodies and apical dendrites [71]. *GRIK3* and *GRIK5* are moderately expressed in the septum from embryonic development, with these levels remaining unchanged in the lateral septum into adulthood [134, 135]. *GRIK2*, *GRIK3*, and *GRIK5* are the only kainate receptor genes consistently detected in the striatum during embryonic development, with *GRIK5* remaining abundant into adulthood [134]. These genes are moderate to abundantly expressed in the thalamus at different stages of prenatal development, while *GRIK3* expression remains strong in the reticular nucleus into adulthood. Finally, in the cerebellum, *GRIK2* and *GRIK5* are the predominant kainate receptor genes expressed in granule cells throughout life, while *GRIK3* is the only kainate receptor gene expressed in stellate/basket cells [135, 134].

In a study of the human fetal cortex, *GRIK3* and *GRIK5* mRNA were far more abundant than those of other kainate receptors throughout the different developmental stages measured (between gestation week 8 and week 20) [138]. This broadly matches with data from the prenatal rat brain, in which *GRIK2*, *GRIK3*, and *GRIK5* are the most widely expressed subunits [134]. Moreover, both the expression of receptor mRNA and receptor binding — suggesting full translation into functional subunits — was found to be greater for kainate receptors during fetal development than for any other ionotropic glutamate receptors, especially during the later gestational periods measured [138]. The expression of kainate receptor transcripts is highly synchronized with that of NMDA receptors at each of the different stages measured. It is likely that these subunits are involved in modulating different processes of neurogenesis during ontogeny, including neuronal migration [138].

Somewhat contrary these findings, AMPA receptor binding from gestational week 16.5 to week 26 has been found to be considerably denser in the hippocampus, entorhinal cortex, temporal cortex, cingulate cortex, putamen, and thalamus than for than for the other ionotropic receptors [139]. NMDA receptor densities were also greater than those of kainate receptors, although these were broadly comparable across most regions and time points, perhaps due to the same synchrony observed in other studies [139, 138]. These discrepancies between mRNA measures and receptor binding densities may be down to delays of the translation into full kainate and NMDA receptor subunits in the regions studied [138].

Some evidence for this delayed development of receptor subunits comes from comparisons of receptor binding in the hippocampi of second trimester fetuses with postmortem measures of full-term infants, three-month-olds, and adults: NMDA receptors made up around half of total glutamate receptors in the infant hippocampi measured, as compared to around a third during the second trimester of gestation or in adults. This was especially the case in the stratum lucidum (CA3) and the molecular layer of the fascia dentata (DG). Moreover, the percentage of the remaining glutamate receptors that bound quisqualate was minimal throughout development [140]. Given that quisqualate binds AMPA receptors with much higher affinity than kainate receptors, a considerable percentage of the remaining glutamate receptors may be expected to be kainate preferring [141, 142, 140]. A separate study of kainate binding in the newborn hippocampus confirms this, with a marked increase relative to prenatal or adult levels in the stratum lucidum and the molecular layer of the fascia dentata, again suggesting synchrony between kainate and NMDA receptor expression during development [143]. Given the relative abundance of *GRIK3* and *GRIK5* expression during earlier stages of human fetal development, it is not implausible that these receptor genes also drive the increase in postnatal expression.

Postmortem mRNA measures taken from adult human brains confirm *GRIK2*, *GRIK3*, and *GRIK5* as the highest expressed kainate receptor genes. However, lower relative expression patterns have been detected for *GRIK3* in adult humans compared to adult rat or fetal human brains. Overall, the dentate gyrus and CA3 remain regions of highest expression, with *GRIK2* and *GRIK5* predominant [66]. It should be noted that studies of adult mRNA expression and receptor binding generally find decreased densities in the human hippocampus relative to rodents and monkeys [144, 145]. However, contrary to the findings of the human study mentioned above [66], it has been posited that these decreases are primarily due to lower expression of *GRIK2* and *GRIK4*, rather than *GRIK3* and *GRIK5* [144]. Moreover, during the postnatal peak of kainate binding in CA3 and DG, human densities of kainate receptors are comparable to or even exceed those of the rat [143]. Possibly, then, selective pressures in our lineage may have driven the increased ontogenetic expression of *GRIK3* and *GRIK5*.

Non-selective immunolabeling of GRIK1-GRIK3 in the monkey hippocampus suggest more qualitative differences in localization as compared with both human and rat data. These low-affinity subunits were found to be densest in

CA1 and the subiculum rather than DG or CA3 [146]. The authors of this study suggest that discrepancies identified between the monkey and the rat may be down to species differences. By comparison, a similar postmortem study in humans found strongest labeling of these subunits in CA2 and CA3 pyramidal neurons (CA2 most abundantly) [147]. Again, it cannot be ruled out that human-monkey differences in low-affinity kainate receptor expression are driven in part by the regulatory changes on *GRIK3* that came under selection in our lineage.

In histological studies of the rat hypothalamus, GRIK2, GRIK3, and GRIK5 receptor proteins have been detected at moderate levels in the supraoptic nucleus (SON), arcuate nucleus, and the junction of the median eminence and infundibulum. Light to moderate staining was detected in the retrochiasmatic region of the SON and the medial mammillary nucleus [71]. Another study found that non-NMDA (mainly kainate) receptors were bound most predominantly in the SON, anterior hypothalamic area, and paraventricular nucleus (PVN) [148]. Both *GRIK2* and *GRIK5* subunit mRNAs are expressed in the anterior parvocellular region of the PVN, while *GRIK5* is highest expressed in the medial parvocellular nucleus and is expressed on CRH-releasing neurons [149]. *GRIK2* and *GRIK3* are most highly expressed in the the subparaventricular zone and perinuclear region surrounding the PVN [150]. Studies *in vitro* confirm that kainate receptors are extensively expressed in the hypothalamus during embryonic development [151], although we are unaware of any study detailing specific subunit abundances or localizations during ontogeny.

These expression data underscore the particular importance of kainate receptor genes that most often show signals of selection in domesticates and modern humans. Evidence that *GRIK3* and *GRIK5* are the kainate receptor subunits with highest expression levels in the developing human brain is suggestive that they are important for neurogenesis and differentiation. Although no easy comparisons can be made between the brain-expression data for embryonic or perinatal rats and a human fetuses, *GRIK3* and *GRIK5* expression levels are high in each species, suggesting that similar developmental processes are regulated by these subunits. The comparatively higher expression of *GRIK2* as compared with *GRIK3* in the rat suggests there may be a more prominent role for the former in CNS development in mammals other than humans. If this is so, human *GRIK3* may occupy homologous ontogenetic functions to *GRIK2*, perhaps helping to explain the tendency for both *GRIK2* and *GRIK3* to show signals of selection in domesticated species, whereas this is only the case for *GRIK3* in AMH.

The relative abundance of kainate receptor gene expression in limbic structures (e.g. hippocampus, entorhinal cortex, cingulate cortex, and, to a lesser extent, the lateral septum and hypothalamus) points to the importance of these genes in the primary functions of these structures, including control of emotions, memory, learning, and neuroendocrine modulation. This is especially the case during the postnatal period, when kainate receptor genes are highest expressed. The cingulate cortex and hippocampus, which prominently modulate HPA-feedback activity, retain moderate to high expression of these genes into adulthood. As detailed above, in the DG and CA3, *GRIK2* and *GRIK5* are expressed most abundantly in the adult rat brain, while *GRIK3* and *GRIK5* are abundant in DG during early postnatal development.

#### Metabotropic glutamate receptor expression

Group II and Group III metabotropic glutamate receptor genes are expressed throughout the central nervous system and are often highly expressed in limbic structures, including the hippocampal formation, as well as in sensory pathways, such as the olfactory and visual systems [73]. The Group II receptor genes, *GRM2* and *GRM3* are moderate to abundantly expressed in various regions of the limbic cortex, Golgi cells of the cerebellum, the dentate gyrus, and in distinct amygdaloid nuclei [152, 153]. Of the Group III receptor genes, *GRM4*, *GRM7*, and *GRM8* are distributed widely across the brain, *GRM7* most expansively and abundantly, followed by *GRM4*, and *GRM8*, which is widely distributed, but expressed in lower quantities [154, 155]. *GRM6* expression is limited to the ON-bipolar cells of the retina. Group II and III receptors play a more prominent role than Group I in the presynaptic modulation of glutamatergic signaling [73, 155].

In the rat brain, *GRM2* is most intensely expressed in the in Golgi cells of the cerebellum, accessory olfactory bulb, and anterior olfactory nucleus, followed by, the entorhinal and parasubicular cortices, granule cells of the dentate gyrus, basolateral and basomedial amygdala, various thalamic nuclei, the medial mammillary nucleus of the hypothalamus, and the cingulate and retrosplenial cortices [152]. Human *GRM2* mRNA is most prominently detected in the cerebellum, hypothalamus, and thalamus, with lighter expression in the hippocampus [156].

In the rat, *GRM3* expression is most abundant in the thalamic reticular nucleus, followed by the lateral and basolateral amygdala, Golgi cells of the cerebellum, the cingulate, retrosplenial, perirhinal, entorhinal, and parasubicular cortices, distinct subparts of the brain stem, and the supraoptic nucleus of the hypothalamus, with lighter expression in the paraventricular and lateral areas of the hypothalamus, as well as granule cells of the dentate gyrus, and stellate/basket cells of the cerebellum [153]. In humans, *GRM3* mRNA has been shown to be similarly expressed in the dentate gyrus, cerebellum, and thalamic reticular nucleus, although there is also evidence of more extensive expression across thalamic nuclei and cerebellar cell types. Strong GRM3 receptor immunoreactivity signals have been detected in the human prefrontal cortex, and low to moderate levels in the hippocampus, amygdala, and thalamus [157].

Non-selective labeling of Group II receptors in the human hippocampus identified dense staining in the dentate gyrus, CA2-CA4, and granule cells of CA3, with lighter staining in CA1 and the subicular cortex [158]. Elsewhere in the brain Group II immunoreactivity is present in the prefrontal cortex and striatum [157]

*GRM4* is highest expressed in the rat sensory ganglia and in granule cells of the cerebellum. In general, it is more extensively expressed than both Group II receptors in the olfactory system and septum, but less so in the limbic cortices.

Expression in the amygdala is limited to the rostral portion of the intercalated nuclei. Like *GRM2*, expression of *GRM4* within the rat hypothalamus is mostly limited to the medial mammillary nucleus. Moderate expression within the hippocampus is mostly limited to the rat dentate gyrus, with similar levels detected in the entorhinal cortex [154]. However, immunocytochemical localization has detected more intense labeling of GRM4 proteins on cell bodies and apical dendrites in CA1 and CA3, and on the bodies of granule cells in DG [159]. In humans, *GRM4* mRNA is strongest detected in the cerebellum, and more moderately in the hippocampus, hypothalamus, and thalamus [160]. In a postmortem human immunohistochemical study of the hippocampus, a specific GRM4 isoform showed moderate labeling on CA3 mossy fibres, with weaker labeling in CA2. Labeling was also detected on the hippocampal efferents of the alveus, parts of which contribute to inputs of the lateral septal nuclei [158, 161]. These data largely parallel with data from the rat, although there is evidence of higher *GRM4* mRNA levels in the human caudate nucleus and putamen, which form part of the dorsal striatum [162].

*GRM7* is expressed in almost all brain regions in the rat, generally at moderate levels. This includes all limbic, amygdalar, hippocampal, and thalamic regions, and practically all hypothalamic subdivisions. Like GRM4, intense immunocytochemical staining can be detected for GRM7 subunits in all regions of the hippocampus, although this is most apparent in neuropil rather than on cell bodies [159]. Highest gene expression is detected in the sensory ganglia, the locus coeruleus within the brainstem, the main olfactory bulb, and the medial septal nucleus [154]. In accordance with studies of the rat, human *GRM7* mRNA levels are also widely distributed at moderate levels in various regions of the human brain, being highest in the neocortex, cerebellum, and regions DG and CA3 of the hippocampus, followed by CA1 [163, 164]. Expression was also detected in the thalamus, with lower levels in the caudate-putamen, in contrast to studies of the rat and the inverse of species expression differences for *GRM4* [164].

In the rat *GRM8* is highest expressed in the olfactory bulb, pontine nuclei of the brainstem, piriform cortex, and reticular nucleus of the thalamus, with lower expression in the the neocortex, hippocampus, basolateral amygdala, and mammillary body of the hypothalamus [165, 166, 163]. In humans, distinct *GRM8* isoforms are differentially but widely expressed, with the most abundant variant showing highest expression in the thalamus, subthalamic nucleus, caudate nucleus, and amygdala, with a second variant being expressed in the visual cortex, caudate-putamen, and cerebellum [167].

Among all glutamate receptor subtypes, metabotropic glutamate receptor binding densities in the rat hypothalamus are second only to NMDA receptors (particularly in dorsomedial, ventromedial, and paraventricular regions) [148]. Given the mRNA expression levels reviewed above and relatively low Group I metabotropic receptor expression in the adult rat hypothalamus, a considerable percentage of these metabotropic receptors may be expected to comprise multiple Group II and III subtypes, most prominently in the medial mammillary nucleus [168, 152, 153, 154]. GRM2 and GRM3 receptor proteins have been detected in the mouse and rat hypothalami and have been shown to modulate HPA function [169].

Unlike kainate receptors, developmental regulation of Group II and III metabotropic receptors in the rat hypothalamus may tend towards increased expression in adulthood. Such increases have been identified in both arcuate and suprachiasmatic nuclei for Group II and III subunits [170]. On the other hand, across brain regions, most metabotropic receptors show similar patterns to kainate receptors, peaking in the early postnatal period. *GRM2* shows a progressive increase in expression during postnatal development in the rat, but remains high in the entorhinal cortex, DG, and various nuclei of the thalamus into adulthood. Low expression levels were found in CA1 and CA3 (in contrast to the human postmortem study described above [158]), and none was detected in the striatum [171]. In the human fetal brain *GRM2* expression is higher than in that of adults, although no regional-specific prenatal expression has been described [156]. *GRM3* generally has highest postnatal expression at birth, especially in the striatum and cerebellum, decreasing progressively throughout development, but remaining high in the thalamic reticular nucleus into adulthood [171, 172].

Expression levels for *GRM4* were found to be low throughout development in one study of the rat, with little change into adulthood, except in granule cells of the cerebellum, where they rise considerably [171]. Other studies have found a similar rise in *GRM4* expression on cell bodies in all major regions of the hippocampus in adulthood [172]. GRM7 proteins show increased postnatal abundance in the rat relative to adult levels across almost all regions studied, including the pons, cerebellum, thalamus, hippocampus, olfactory cortex, and striatum, although expression remains relatively high in the hippocampus, bed nucleus of stria terminalis (BNST), medial mammillary body of the hypothalamus, and various thalamic nuclei into adulthood [173]. In humans, *GRM8* shows increased expression in the fetal brain than in that of adults [167].

Kainate and Group II and III metabotropic receptors have broadly overlapping localizations and timings of expression, with predominant postnatal expression in limbic cortices, the hippocampus (especially DG and CA3), cerebellum, thalamus (particularly the reticular nucleus), and hypothalamus (most notably the medial mammillary nucleus, although perhaps asynchronously). Differences include higher metabotropic expression in sensory pathways, as well as the amygdala, thalamus, distinct hippocampal regions, and in the adult hypothalamus more generally. Kainate receptor expression seems to be more predominant in the cingulate cortex and striatum, particularly during ontogeny. Both receptor subtypes have metabotropic functions modulating the release of glutamate, and both are part of G-protein coupled signaling cascades. Together, these receptor subtypes are extensively expressed in regions crucial for the modualtion of how an organism deals with stress, from the moment sensory inputs for a stressor are received right through to regulation of feedback mechanisms that inform best responses.

### S4 Glutamatergic signaling-Neural Crest interactions

In this section we discuss two ways in which glutamatergic signaling may interact with, or even bypass, the neural crest to bring about the broader phenotypical features of the domestication syndrome.

Under one account, the reduction in neural crest cell (NCC) proliferation may occur subsequent to changes in glutamatergic transmission. Modulation of glutamatergic signaling attenuates HPA-axis signaling, and ultimately glucocorticoid output, from the adrenal cortex in (pre)domesticated females during pregnancy. Modified hormonal concentrations would then affect embryonic development, altering neural crest cell inputs to different tissues prenatally. In cell cultures, glucocorticoids are essential for NCC survival and differentiation [174]. An increase in corticosteroid concentrations has been shown to increase NCC numbers and alter cell fates, for example converting small intensely fluorescent cells, which are normally dopaminergic, into epinephrine-synthesizing cells. Removal of corticosteroids and replacement with nerve growth factor converts small intensely fluorescent cells and chromaffin cells into sympathetic neurons [175, 174]. Mice embryos lacking the glucocorticoid receptor cannot survive outside the womb due to widespread defects, including severe reductions in chromaffin cells from early development. Those cells that remain lose their ability to synthesize epinephrine [176]. The above studies suggest that lowered stress hormone levels in the womb, resulting from HPA hypofunction in the mother can have the knock-on effect of bringing about neural crest hypofunction and cell-differentiation changes in embryonic development. Under our hypothesis, mild reductions in NCC inputs may be driven by increased glutamatergic modulation of the stress response during pregnancy resulting from selection for tameness.

As mentioned in the main text, corticosterone treatment during embryonic development of domesticated ducks can bring about mallard-like behaviors in neonates [177, 178]. Glucocorticoid levels are lower during gestation in both tame rats and foxes compared with aggressive strains [179]. Suppression of glucocorticoid levels in pregnant aggressive females between days 12 and 14 of gestation leads to a concomitant reduction in embryonic glucocorticoids by day 20. This, in turn, leads to significant increases in depigmentation in neonate rats [179]. In both the rat and duck studies, the effects of glucocorticoids on embryonic development take place long after neurulation and migration of neural crest cells away from the neural tube. This suggests that at least some of the most important phenotypical features of domestication need not depend on genetic changes to NCC development or migration, but, rather, on postmigratory regulation of expression at different tissue sites by glucocorticoids.

An alternative possibility is that alterations to glutamate receptor signaling can directly account for both behavioral and physical changes in domestication. Some evidence for this derives from the fact that glutamate receptor, transporter, and other signaling genes identified in different domestication studies are expressed in melanocytes, osteoblasts, osteoclasts, chondrocytes, and other tissues beyond the CNS [180, 181, 182, 183, 184]. Thus, traits such as floppy ears, shortened snouts, or depigmentation could feasibly result from alterations to glutamatergic receptor expression in these tissues. Bone is innervated by glutamatergic fibers and glutamate receptors are thought to play a role in osteoblastic differentiation and proliferation [185, 186]. *GRIK5*, *GRM4*, and *GRM8* along with NMDA and AMPA receptor genes are expressed *in vitro* in developing rat calvarial osteoblasts [187, 188]. Group II and III metabotropic receptors have been shown to prevent mineralization of chondrocytes, implicating them in cartilage development [180, 184]. NMDA receptors have been implicated in the maturation and differentiation of chondrocytes [189]. Glutamate receptor genes (among them *GRIK3*) are expressed on melanocytes, and inhibition of AMPA-mediated excitatory transmission has been found to decrease the expression of *MITF* (melanogenesis associated transcription factor) [181].

[6] Stéphane Peyrégne, Michael James Boyle, Michael Dannemann, and Kay

Prüfer. Detecting ancient positive selection in humans using extended lineage sorting. *Genome Research*, 27(9):1563–1572, September 2017.

[7] Grégory Gauvain, Ingrid Chamma, Quentin Chevy, Carolina Cabezas, Theano Irinopoulou, Natalia Bodrug, Michèle Carnaud, Sabine Lévi, and Jean Christophe Poncer. The neuronal K-Cl cotransporter KCC2 influences postsynaptic AMPA receptor content and lateral diffusion in dendritic spines. *Proceedings of the National Academy of Sciences*, 108(37):15474–15479, September 2011.

[8] Evgueni A. Ivakine, Brooke A. Acton, Vivek Mahadevan, Jake Ormond, Man Tang, Jessica C. Pressey, Michelle Y. Huang, David Ng, Eric Delpire, Michael W. Salter, Melanie A. Woodin, and Roderick R. McInnes. Neto2 is a KCC2 interacting protein required for neuronal Cl- regulation in hippocampal neurons. *Proceedings of the National Academy of Sciences*, page 201212907, February 2013.

[9] Ran Tao, Chao Li, Erin N. Newburn, Tianzhang Ye, Barbara K. Lipska, Mary M. Herman, Daniel R. Weinberger, Joel E. Kleinman, and Thomas M. Hyde. Transcript-Specific Associations of SLC12a5 (KCC2) in Human Prefrontal Cortex with Development, Schizophrenia, and Affective Disorders. *Journal of Neuroscience*, 32(15):5216–5222, April 2012.

[10] Nancy D. Merner, Madison R. Chandler, Cynthia Bourassa, Bo Liang, Arjun R. Khanna, Patrick Dion, Guy A. Rouleau, and Kristopher T. Kahle. Regulatory domain or CpG site variation in SLC12a5, encoding the chloride transporter KCC2, in human autism and schizophrenia. *Frontiers in Cellular Neuroscience*, 9, 2015.

[11] Tommy Stödberg, Amy McTague, Arnaud J. Ruiz, Hiromi Hirata, Juan Zhen, Philip Long, Irene Farabella, Esther Meyer, Atsuo Kawahara, Grace Vassallo, Stavros M. Stivaros, Magnus K. Bjursell, Henrik Stranneheim, Stephanie Tigerschiöld, Bengt Persson, Iftikhar Bangash, Krishna Das, Deborah Hughes, Nicole Lesko, Joakim Lundeberg, Rod C. Scott, Annapurna Poduri, Ingrid E. Scheffer, Holly Smith, Paul Gissen, Stephanie Schorge, Maarten E. A. Reith, Maya Topf, Dimitri M. Kullmann, Robert J. Harvey, Anna Wedell, and Manju A. Kurian. Mutations in *SLC12A5* in epilepsy of infancy with migrating focal seizures. *Nature Communications*, 6:8038, September 2015.

[12] Hirotomo Saitsu, Miho Watanabe, Tenpei Akita, Chihiro Ohba, Kenji Sugai, Winnie Peitee Ong, Hideaki Shiraishi, Shota Yuasa, Hiroshi Matsumoto, Khoo Teik Beng, Shinji Saitoh, Satoko Miyatake, Mitsuko Nakashima, Noriko Miyake, Mitsuhiro Kato, Atsuo Fukuda, and Naomichi Matsumoto. Impaired neuronal KCC2 function by biallelic *SLC12A5* mutations in migrating focal seizures and severe developmental delay. *Scientific Reports*, 6:30072, July 2016.

[13] Florian J. Alberto, Frédéric Boyer, Pablo Orozco-terWengel, Ian Streeter, Bertrand Servin, Pierre Villemereuil, Badr Benjelloun, Pablo Librado, Filippo Biscarini, Licia Colli, Mario Barbato, Wahid Zamani, Adriana Alberti, Stefan Engelen, Alessandra Stella, Stéphane Joost, Paolo Ajmone-Marsan, Riccardo Negrini, Ludovic Orlando, Hamid Reza Rezaei, Saeid Naderi, Laura Clarke, Paul Flicek, Patrick Wincker, Eric Coissac, James Kijas, Gwenola Tosser-Klopp, Abdelkader Chikhi, Michael W. Bruford, Pierre Taberlet, and François Pompanon. Convergent genomic signatures of domestication in sheep and goats. *Nature Communications*, 9(1):813, March 2018.

[14] Henrike O. Heyne, Susann Lautenschläger, Ronald Nelson, François Besnier, Maxime Rotival, Alexander Cagan, Rimma Kozhemyakina, Irina Z. Plyusnina, Lyudmila Trut, Örjan Carlborg, Enrico Petretto, Leonid Kruglyak, Svante Pääbo, Torsten Schöneberg, and Frank W. Albert. Genetic Influences on Brain Gene Expression in Rats Selected for Tameness and Aggression. *Genetics*, 198(3):1277–1290, November 2014.

[15] Swapan Mallick, Heng Li, Mark Lipson, Iain Mathieson, Melissa Gymrek, Fernando Racimo, Mengyao Zhao, Niru Chennagiri, Susanne Nordenfelt, Arti Tandon, Pontus Skoglund, Iosif Lazaridis, Sriram Sankararaman, Qiaomei Fu, Nadin Rohland, Gabriel Renaud, Yaniv Erlich, Thomas Willems, Carla Gallo, Jeffrey P. Spence, Yun S. Song, Giovanni Poletti, Francois Balloux, George van Driem, Peter de Knijff, Irene Gallego Romero, Aashish R. Jha, Doron M. Behar, Claudio M. Bravi, Cristian Capelli, Tor Hervig, Andres Moreno-Estrada, Olga L. Posukh, Elena Balanovska, Oleg Balanovsky, Sena Karachanak-Yankova, Hovhannes Sahakyan, Draga Toncheva, Levon Yepiskoposyan, Chris Tyler-Smith, Yali Xue, M. Syafiq Abdullah, Andres Ruiz-Linares, Cynthia M. Beall, Anna Di Rienzo, Choongwon Jeong, Elena B. Starikovskaya, Ene Metspalu, Jüri Parik, Richard Villems, Brenna M. Henn, Ugur Hodoglugil, Robert Mahley, Antti Sajantila, George Stamatoyannopoulos, Joseph T. S. Wee, Rita Khusainova, Elza Khusnutdinova, Sergey Litvinov, George Ayodo, David Comas, Michael F. Hammer, Toomas Kivisild, William Klitz, Cheryl A. Winkler, Damian Labuda, Michael Bamshad, Lynn B. Jorde, Sarah A. Tishkoff, W. Scott Watkins, Mait Metspalu, Stanislav Dryomov, Rem Sukernik, Lalji Singh, Kumarasamy Thangaraj, Svante Pääbo, Janet Kelso, Nick Patterson, and David Reich. The Simons Genome Diversity Project: 300 genomes from 142 diverse populations. *Nature*, 538(7624):201–206, October 2016.

[16] S. E. Stewart, D. Yu, J. M. Scharf, B. M. Neale, J. A. Fagerness, C. A. Mathews, P. D. Arnold, P. D. Evans, E. R. Gamazon, L. Osiecki, L. McGrath, S. Haddad, J. Crane, D. Hezel, C. Illman, C. Mayerfeld, A. Konkashbaev, C. Liu, A. Pluzhnikov, A. Tikhomirov, C. K. Edlund, S. L. Rauch, R. Moessner, P. Falkai, W. Maier, S. Ruhrmann, H.-J. Grabe, L. Lennertz, M. Wagner, L. Bellodi, M. C. Cavallini, M. A.

Richter, E. H. Cook Jr, J. L. Kennedy, D. Rosenberg, D. J. Stein, S. M. J. Hemmings, C. Lochner, A. Azzam, D. A. Chavira, E. Fournier, H. Garrido, B. Sheppard, P. Umaña, D. L. Murphy, J. R. Wendland, J. Veenstra-VanderWeele, D. Denys, R. Blom, D. Deforce, F. Van Nieuwerburgh, H. G. M. Westenberg, S. Walitza, K. Egberts, T. Renner, E. C. Miguel, C. Cappi, A. G. Hounie, M. Conceição do Rosário, A. S. Sampaio, H. Vallada, H. Nicolini, N. Lanzagorta, B. Camarena, R. Delorme, M. Leboyer, C. N. Pato, M. T. Pato, E. Voyiaziakis, P. Heutink, D. C. Cath, D. Posthuma, J. H. Smit, J. Samuels, O. J. Bienvenu, B. Cullen, A. J. Fyer, M. A. Grados, B. D. Greenberg, J. T. McCracken, M. A. Riddle, Y. Wang, V. Coric, J. F. Leckman, M. Bloch, C. Pittenger, V. Eapen, D. W. Black, R. A. Ophoff, E. Strengman, D. Cusi, M. Turiel, F. Frau, F. Macciardi, J. R. Gibbs, M. R. Cookson, A. Singleton, North American Brain Expression Consortium, S. Arepalli, M. R. Cookson, A. Dillman, L. Ferrucci, J. R. Gibbs, D. G. Hernandez, R. Johnson, D. L. Longo, M. A. Nalls, R. O'Brien, A. Singleton, B. Traynor, J. Troncoso, M. van der Brug, H. R. Zielke, A. Zonderman, J. Hardy, UK Brain Expression Database, J. A. Hardy, M. Ryten, C. Smith, D. Trabzuni, R. Walker, Mike Weale, A. T. Crenshaw, M. A. Parkin, D. B. Mirel, D. V. Conti, S. Purcell, G. Nestadt, G. L. Hanna, M. A. Jenike, J. A. Knowles, N. Cox, and D. L. Pauls. Genome-wide association study of obsessive-compulsive disorder. *Molecular Psychiatry*, 18(7):788–798, July 2013.

[17] Jiang Li, Jiajia Cui, Xiuhai Wang, Jianhua Ma, Haitao Niu, Xu Ma, Xinhua Zhang, and Shiguo Liu. An association study between DLGAP1 rs11081062 and EFNA5 rs26728 polymorphisms with obsessive–compulsive disorder in a Chinese Han population. *Neuropsychiatric Disease and Treatment*, 11:897–905, April 2015.

[18] M. Mattheisen, J. F. Samuels, Y. Wang, B. D. Greenberg, A. J. Fyer, J. T. McCracken, D. A. Geller, D. L. Murphy, J. A. Knowles, M. A. Grados, M. A. Riddle, S. A. Rasmussen, N. C. McLaughlin, E. L. Nurmi, K. D. Askland, H.-D. Qin, B. A. Cullen, J. Piacentini, D. L. Pauls, O. J. Bienvenu, S. E. Stewart, K.-Y. Liang, F. S. Goes, B. Maher, A. E. Pulver, Y. Y. Shugart, D. Valle, C. Lange, and G. Nestadt. Genome-wide association study in obsessive-compulsive disorder: results from the OCGAS. *Molecular Psychiatry*, 20(3):337–344, March 2015.

[19] Mikkel Schubert, Hákon Jónsson, Dan Chang, Clio Der Sarkissian, Luca Ermini, Aurélien Ginolhac, Anders Albrechtsen, Isabelle Dupanloup, Adrien Foucal, Bent Petersen, Matteo Fumagalli, Maanasa Raghavan, Andaine Seguin-Orlando, Thorfinn S. Korneliussen, Amhed M. V. Velazquez, Jesper Stenderup, Cindi A. Hoover, Carl-Johan Rubin, Ahmed H. Alfarhan, Saleh A. Alquraishi, Khaled A. S. Al-Rasheid, David E. MacHugh, Ted Kalbfleisch, James N. MacLeod, Edward M. Rubin, Thomas Sicheritz-Ponten, Leif Andersson, Michael Hofreiter, Tomas Marques-Bonet, M. Thomas P. Gilbert, Rasmus Nielsen, Laurent Ex-

coffier, Eske Willerslev, Beth Shapiro, and Ludovic Orlando. Prehistoric genomes reveal the genetic foundation and cost of horse domestication. *Proceedings of the National Academy of Sciences*, 111(52):E5661–E5669, December 2014.

[28] Anna V. Kukekova, Jennifer L. Johnson, Xueyan Xiang, Shaohong Feng, Shiping Liu, Halie M. Rando, Anastasiya V. Kharlamova, Yury Herbeck, Natalya A. Serdyukova, Zijun Xiong, Violetta Beklemischeva, Klaus-Peter Koepfli, Rimma G. Gulevich, Anastasiya V. Vladimirova, Jessica P. Hekman, Polina L. Perelman, Aleksander S. Graphodatsky, Stephen J.

O'Brien, Xu Wang, Andrew G. Clark, Gregory M. Acland, Lyudmila N. Trut, and Guojie Zhang. Red fox genome assembly identifies genomic regions associated with tame and aggressive behaviours. *Nature Ecology & Evolution*, 2(9):1479–1491, September 2018.

[29] Jeffrey N. Savas, Luís F. Ribeiro, Keimpe D. Wierda, Rebecca Wright, Laura A. DeNardo-Wilke, Heather C. Rice, Ingrid Chamma, Yi-Zhi Wang, Roland Zemla, Mathieu Lavallée-Adam, Kristel M. Vennekens, Matthew L. O'Sullivan, Joseph K. Antonios, Elizabeth A. Hall, Olivier Thoumine, Alan D. Attie, John R. Yates, Anirvan Ghosh, and Joris de Wit. The Sorting Receptor SorCS1 Regulates Trafficking of Neurexin and AMPA Receptors. *Neuron*, 87(4):764–780, August 2015.

[30] Stéphane Peyrégne, Michael James Boyle, Michael Dannemann, and Kay Pruefer. Detecting ancient positive selection in humans using extended lineage sorting. *bioRxiv*, December 2016.

[48] Elena A. B. Azizan, Hanne Poulsen, Petronel Tuluc, Junhua Zhou, Michael V. Clausen, Andreas Lieb, Carmela Maniero, Sumedha Garg, Elena G. Bochukova, Wanfeng Zhao, Lalarukh Haris Shaikh, Cheryl A. Brighton, Ada E. D. Teo, Anthony P. Davenport, Tanja Dekkers, Bas Tops, Benno Küsters, Jiri Ceral, Giles S. H. Yeo, Sudeshna Guha Neogi, Ian McFarlane, Nitzan Rosenfeld, Francesco Marass, James Hadfield, Wojciech Margas, Kanchan Chaggar, Miroslav Solar, Jaap Deinum, Annette C. Dolphin, I. Sadaf Farooqi, Joerg Striessnig, Poul Nissen, and Morris J. Brown. Somatic mutations in *ATP1A1* and *CACNA1D* underlie a common subtype of adrenal hypertension. *Nature Genetics*, 45(9):1055–1060, September 2013.

[49] Mary Ann Raghanti, Melissa K. Edler, Alexa R. Stephenson, Emily L. Munger, Bob Jacobs, Patrick R. Hof, Chet C. Sherwood, Ralph L. Holloway, and C. Owen Lovejoy. A neurochemical hypothesis for the origin of hominids. *Proceedings of the National Academy of Sciences*, page 201719666, January 2018.

[50] François Beaubien, Reesha Raja, Timothy E. Kennedy, Alyson E. Fournier, and Jean-François Cloutier. Slitrk1 is localized to excitatory synapses and promotes their development. *Scientific Reports*, 6:27343, June 2016.

[51] Martin Kuhlwilm and Cedric Boeckx. Genetic differences between humans and other hominins contribute to the "human condition". *bioRxiv*, April 2018.

[52] Jesse F. Abelson, Kenneth Y. Kwan, Brian J. O'Roak, Danielle Y. Baek, Althea A. Stillman, Thomas M. Morgan, Carol A. Mathews, David L. Pauls, Mladen-Roko Rašin, Murat Gunel, Nicole R. Davis, A. Gulhan Ercan-Sencicek, Danielle H. Guez, John A. Spertus, James F. Leckman, Leon S. Dure, Roger Kurlan, Harvey S. Singer, Donald L. Gilbert, Anita Farhi, Angeliki Louvi, Richard P. Lifton, Nenad Šestan, and Matthew W. State. Sequence Variants in SLITRK1 Are Associated with Tourette's Syndrome. *Science*, 310(5746):317–320, October 2005.

[53] Yun Zhou and Niels Christian Danbolt. GABA and Glutamate Transporters in Brain. *Frontiers in Endocrinology*, 4(165), November 2013.

[54] Paul Daniel Arnold, Tricia Sicard, Eliza Burroughs, Margaret A. Richter, and James L. Kennedy. Glutamate Transporter Gene SLC1a1 Associated With Obsessive-compulsive Disorder. *Archives of General Psychiatry*, 63(7):769–776, July 2006.

[55] Rageen Rajendram, Sefi Kronenberg, Christie L. Burton, and Paul D. Arnold. Glutamate Genetics in Obsessive-Compulsive Disorder: A Review. *Journal of the Canadian Academy of Child and Adolescent Psychiatry*, 26(3):205–213, 2017.

[56] P. G. Eusebi, O. Cortés, C. Carleos, S. Dunner, and J. Cañon. Detection of selection signatures for agonistic behaviour in cattle. *Journal of Animal Breeding and Genetics = Zeitschrift Fur Tierzuchtung Und Zuchtungsbiologie*, April 2018.

[89] Mònica Gratacòs, Javier Costas, Rafael de Cid, Mònica Bayés, Juan R. González, Enrique Baca-García, Yolanda de Diego, Fernando Fernández-Aranda, José Fernández-Piqueras, Miriam Guitart, Rocío Martín-Santos, Lourdes Martorell, José M. Menchón, Miquel Roca, Jerónimo Sáiz-Ruiz, Julio Sanjuán, Marta Torrens, Mikel Urretavizcaya, Joaquín Valero, Elisabet Vilella, Xavier Estivill, and Ángel Carracedo. Identification of new putative susceptibility genes for several psychiatric disorders by association analysis of regulatory and non-synonymous SNPs of 306 genes involved in neurotransmission and neurodevelopment. *American Journal of Medical Genetics Part B: Neuropsychiatric Genetics*, 150B(6):808–816, September 2009.

[90] Niklas Krumm, Tychele N. Turner, Carl Baker, Laura Vives, Kiana Mohajeri, Kali Witherspoon, Archana Raja, Bradley P. Coe, Holly A. Stessman, Zong-Xiao He, Suzanne M. Leal, Raphael Bernier, and Evan E. Eichler. Excess of rare, inherited truncating mutations in autism. *Nature Genetics*, 47(6):582–588, June 2015.

[91] Xu Wang, Lenore Pipes, Lyudmila N. Trut, Yury Herbeck, Anastasiya V. Vladimirova, Rimma G. Gulevich, Anastasiya V. Kharlamova, Jennifer L. Johnson, Gregory M. Acland, Anna V. Kukekova, and Andrew G. Clark. Genomic responses to selection for tame/aggressive behaviors in the silver fox (Vulpes vulpes). *Proceedings of the National Academy of Sciences*, page 201800889, September 2018.

[99] Niamh L. O'Brien, Michael J. Way, Radhika Kandaswamy, Alessia Fiorentino, Sally I. Sharp, Giorgia Quadri, Jarram Alex, Adebayo Anjorin, David Ball, Raquin Cherian, Karim Dar, Aynur Gormez, Irene Guerrini, Mathis Heydtmann, Audrey Hillman, Sudheer Lankappa, Greg Lydall, Aideen O'Kane, Shamir Patel, Digby Quested, Iain Smith, Allan D. Thomson, Nicholas J. Bass, Marsha Y. Morgan, David Curtis, and Andrew McQuillin. The functional GRM3 Kozak sequence variant rs148754219 affects the risk of schizophrenia and alcohol dependence as well as bipolar disorder:. *Psychiatric Genetics*, 24(6):277–278, December 2014.

[100] Jillian P. Casey, Tiago Magalhaes, Judith M. Conroy, Regina Regan, Naisha Shah, Richard Anney, Denis C. Shields, Brett S. Abrahams, Joana Almeida, Elena Bacchelli, Anthony J. Bailey, Gillian Baird, Agatino Battaglia, Tom Berney, Nadia Bolshakova, Patrick F. Bolton, Thomas Bourgeron, Sean Brennan, Phil Cali, Catarina Correia, Christina Corsello, Marc Coutanche, Geraldine Dawson, Maretha de Jonge, Richard Delorme, Eftichia Duketis, Frederico Duque, Annette Estes, Penny Farrar, Bridget A. Fernandez, Susan E. Folstein, Suzanne Foley, Eric Fombonne, Christine M. Freitag, John Gilbert, Christopher Gillberg, Joseph T. Glessner, Jonathan Green, Stephen J. Guter, Hakon Hakonarson, Richard Holt, Gillian Hughes, Vanessa Hus, Roberta Igliozzi, Cecilia Kim, Sabine M. Klauck, Alexander Kolevzon, Janine A. Lamb, Marion Leboyer, Ann Le Couteur, Bennett L. Leventhal, Catherine Lord, Sabata C. Lund, Elena Maestrini, Carine Mantoulan, Christian R. Marshall, Helen McConachie, Christopher J. McDougle, Jane McGrath, William M. McMahon, Alison Merikangas, Judith Miller, Fiorella Minopoli, Ghazala K. Mirza, Jeff Munson, Stanley F. Nelson, Gudrun Nygren, Guiomar Oliveira, Alistair T. Pagnamenta, Katerina Papanikolaou, Jeremy R. Parr, Barbara Parrini, Andrew Pickles, Dalila Pinto, Joseph Piven, David J. Posey,

Annemarie Poustka, Fritz Poustka, Jiannis Ragoussis, Bernadette Roge, Michael L. Rutter, Ana F. Sequeira, Latha Soorya, Inês Sousa, Nuala Sykes, Vera Stoppioni, Raffaella Tancredi, Maïté Tauber, Ann P. Thompson, Susanne Thomson, John Tsiantis, Herman Van Engeland, John B. Vincent, Fred Volkmar, Jacob A. S. Vorstman, Simon Wallace, Kai Wang, Thomas H. Wassink, Kathy White, Kirsty Wing, Kerstin Wittemeyer, Brian L. Yaspan, Lonnie Zwaigenbaum, Catalina Betancur, Joseph D. Buxbaum, Rita M. Cantor, Edwin H. Cook, Hilary Coon, Michael L. Cuccaro, Daniel H. Geschwind, Jonathan L. Haines, Joachim Hallmayer, Anthony P. Monaco, John I. Nurnberger, Margaret A. Pericak-Vance, Gerard D. Schellenberg, Stephen W. Scherer, James S. Sutcliffe, Peter Szatmari, Veronica J. Vieland, Ellen M. Wijsman, Andrew Green, Michael Gill, Louise Gallagher, Astrid Vicente, and Sean Ennis. A novel approach of homozygous haplotype sharing identifies candidate genes in autism spectrum disorder. *Human Genetics*, 131(4):565–579, April 2012.

[101] Yu-Wen Chen, Hui-Ching Lin, Ming-Chong Ng, Ya-Hsin Hsiao, Chao-Chuan Wang, Po-Wu Gean, and Po See Chen. Activation of mGluR2/3 underlies the effects of N-acetylcystein on amygdala-associated autism-like phenotypes in a valproate-induced rat model of autism. *Frontiers in Behavioral Neuroscience*, 8, 2014.

[102] Lena Wischhof and Michael Koch. Pre-treatment with the mGlu2/3 receptor agonist LY379268 attenuates DOI-induced impulsive responding and regional c-Fos protein expression. *Psychopharmacology*, 219(2):387–400, January 2012.

[103] Andrea Benazzo, Emiliano Trucchi, James A. Cahill, Pierpaolo Maisano Delser, Stefano Mona, Matteo Fumagalli, Lynsey Bunnefeld, Luca Cornetti, Silvia Ghirotto, Matteo Girardi, Lino Ometto, Alex Panziera, Omar Rota-Stabelli, Enrico Zanetti, Alexandros Karamanlidis, Claudio Groff, Ladislav Paule, Leonardo Gentile, Carles Vilà, Saverio Vicario, Luigi Boitani, Ludovic Orlando, Silvia Fuselli, Cristiano Vernesi, Beth Shapiro, Paolo Ciucci, and Giorgio Bertorelle. Survival and divergence in a small group: The extraordinary genomic history of the endangered Apennine brown bear stragglers. *Proceedings of the National Academy of Sciences*, 114(45):E9589–E9597, November 2017.

[114] Clarissa Ganda, Sibylle G. Schwab, Nurmiati Amir, Heriani Heriani, Irman Irmansyah, Agung Kusumawardhani, Martina Nasrun, Ika Widyawati, Wolfgang Maier, and Dieter B. Wildenauer. A family-based association study of DNA sequence variants in GRM7 with schizophrenia

in an Indonesian population. *International Journal of Neuropsychopharmacology*, 12(9):1283–1289, October 2009.

[115] Subin Park, Sun-Woo Jung, Boong-Nyun Kim, Soo-Churl Cho, Min-Sup Shin, Jae-Won Kim, Hee Jeong Yoo, Dae-Yeon Cho, Un-Sun Chung, Jung-Woo Son, and Hyo-Won Kim. Association between the GRM7 rs3792452 polymorphism and attention deficit hyperacitiveity disorder in a Korean sample. *Behavioral and Brain Functions*, 9(1):1, January 2013.

[116] Josephine Elia, Joseph T. Glessner, Kai Wang, Nagahide Takahashi, Corina J. Shtir, Dexter Hadley, Patrick M. A. Sleiman, Haitao Zhang, Cecilia E. Kim, Reid Robison, Gholson J. Lyon, James H. Flory, Jonathan P. Bradfield, Marcin Imielinski, Cuiping Hou, Edward C. Frackelton, Rosetta M. Chiavacci, Takeshi Sakurai, Cara Rabin, Frank A. Middleton, Kelly A. Thomas, Maria Garris, Frank Mentch, Christine M. Freitag, Hans-Christoph Steinhausen, Alexandre A. Todorov, Andreas Reif, Aribert Rothenberger, Barbara Franke, Eric O. Mick, Herbert Roeyers, Jan Buitelaar, Klaus-Peter Lesch, Tobias Banaschewski, Richard P. Ebstein, Fernando Mulas, Robert D. Oades, Joseph Sergeant, Edmund Sonuga-Barke, Tobias J. Renner, Marcel Romanos, Jasmin Romanos, Andreas Warnke, Susanne Walitza, Jobst Meyer, Haukur Pálmason, Christiane Seitz, Sandra K. Loo, Susan L. Smalley, Joseph Biederman, Lindsey Kent, Philip Asherson, Richard J. L. Anney, J. William Gaynor, Philip Shaw, Marcella Devoto, Peter S. White, Struan F. A. Grant, Joseph D. Buxbaum, Judith L. Rapoport, Nigel M. Williams, Stanley F. Nelson, Stephen V. Faraone, and Hakon Hakonarson. Genome-wide copy number variation study associates metabotropic glutamate receptor gene networks with attention deficit hyperactivity disorder. *Nature Genetics*, 44(1):78–84, January 2012.

[130] José V. Negrete-Díaz, Talvinder S. Sihra, Gonzalo Flores, and Antonio Rodríguez-Moreno. Non-canonical Mechanisms of Presynaptic Kainate Receptors Controlling Glutamate Release. *Frontiers in Molecular Neuroscience*, 11, April 2018.

[183] Marcela Julio-Pieper, Peter J. Flor, Timothy G. Dinan, and John F. Cryan. Exciting Times beyond the Brain: Metabotropic Glutamate Receptors in Peripheral and Non-Neural Tissues. *Pharmacological Reviews*, January 2011.
